# Matrix viscoelasticity controls epithelial cell mechanobiology through dimensionality

**DOI:** 10.1101/2024.03.05.583526

**Authors:** Giuseppe Ciccone, Mariana Azevedo Gonzalez Oliva, Marie Versaevel, Marco Cantini, Massimo Vassalli, Manuel Salmeron-Sanchez, Sylvain Gabriele

## Abstract

In recent years, matrix viscoelasticity has emerged as a potent regulator of fundamental cellular processes and has been implicated in promoting cancer progression. Alongside viscoelasticity, additional ECM cues have been shown to influence migration decision-making of cancer cells, and spatial confinement is now considered as a potential regulator of metastasis. However, our understanding of these complex processes predominantly relies on purely elastic hydrogels, and the exact relationship between matrix viscoelasticity and spatial confinement in driving epithelial cell mechanotransduction and migration during cancer progression remains unclear. Here, we systematically investigated the interplay between matrix stiffness, viscoelasticity and spatial confinement by engineering soft (∼0.3 kPa) and stiff (∼3 kPa) polyacrylamide hydrogels with varying degrees of viscous dissipation, mirroring the mechanical properties of healthy and tumoral conditions in breast tissue. We observed that viscoelasticity modulates cell spreading, focal adhesions and YAP nuclear import in opposite directions on soft and stiff substrates. Strikingly, viscoelasticity enhances migration speed and persistence on soft substrates, while impeding them on stiff substrates via actin retrograde flow regulation. Combining soft micropatterning with viscoelastic hydrogels, we also show that spatial confinement restricts cell migration on soft matrices regardless of matrix viscoelasticity and promotes migration on stiff matrices in a viscoelasticity-dependent fashion. Our findings establish substrate viscoelasticity as a key regulator of epithelial cell functions and unravel the role of the matrix dimensionality in this process.

**Significance:** While matrix elasticity has received significant attention, recent findings underscore the importance of its natural dissipative properties and spatial confinement in regulating cellular processes and tumour invasiveness. However, the intricate interplay between viscoelasticity and spatial confinement in orchestrating epithelial cell behaviour during cancer progression remains elusive. Using micropatterned viscoelastic hydrogels to replicate the mechanical properties encountered during breast tumour progression, we unveil that viscoelasticity modulates cell behaviour and mechanotransduction signals differently on soft and stiff substrates. Increased viscoelasticity enhances migration speed and persistence on soft substrates while impeding them on stiff substrates via actin retrograde flow regulation. Furthermore, spatial confinement restricts cell migration on soft matrices regardless of viscoelasticity, while promoting migration on stiff matrices in a viscoelasticity-dependent manner.

## Introduction

The mechanical properties of the extracellular matrix (ECM) have emerged as crucial determinants of cellular behaviour, and changes in ECM mechanics have been linked to disease progression (1, 2). As part of pathophysiological processes, cells actively probe ECM mechanical properties through integrins and associated focal adhesions, forming a dynamic crosstalk that profoundly influences cells spreading, migration and differentiation (3).

ECM mechanical properties have been predominantly replicated in terms of elasticity, commonly referred to as stiffness or rigidity. It is now widely acknowledged that matrix elasticity alone influences numerous pathophysiologically-relevant cellular processes (4). For instance, recent studies have demonstrated that durotaxis, the process of migration along stiffness gradients (5–7), occurs in vivo in the *Xenopus laevis* neural crest, orchestrating the coordinated movement of cells (8).

However, while much attention has been focused on matrix elasticity, ECM mechanics are intrinsically governed by viscoelasticity. Indeed, the mechanical response that cells probe is time dependent, with stress relaxation dictating the perceived mechanical properties that, in turn, drive cell behaviour (9). The ECM is a complex polymeric network embedded in the extracellular fluid, rendering it a viscoelastic material (10, 11). More specifically, ECM viscoelasticity has been associated to the breaking of weak crosslinks, polymer entanglements and protein unfolding (10, 11), resulting in the dissipation of stress exerted by cells over time.

While exploration of relevant relaxation times in vivo is still at its early stages (9, 12), the role of viscoelasticity in influencing cell responses in vitro has gained appreciation over the past decade. Notably, research has shown that matrix energy dissipation plays a regulatory role in fundamental processes such as cell spreading (13–15), differentiation (14, 16–18) and more recently cell migration (19, 20) and cancer progression (12). In the body, cell migration plays a crucial role in diverse processes, including development, wound healing and cancer metastasis (21). The coordinated movement of cells is orchestrated by numerous biophysical factors, such as ECM stiffness (5–7) and spatial confinement imposed by the ECM or neighbouring cells (22–26). Of note, spatial confinement has been shown to influence migration decision-making of cancer cells (27) and is now considered as a potential regulator of metastasis (28).

In fact, cells in the body oftentimes migrate in confined spaces, such as along ECM fibres or through dense tissues, leveraging these guidance cues to facilitate migration along predetermined tracks, a phenomenon notably observed in cancer progression (23). Interestingly, it has been demonstrated that matrix confinement alters the relationship between cell migration speed and ECM stiffness (24, 29, 30). These results indicate a complex interplay between the mechanical properties of the matrix and the level of cellular confinement, both of which play a pivotal role in tumour invasiveness (23, 31). More recently, it has been reported that cancer tissue behaves as a viscoelastic material, exhibiting distinct viscoelastic properties that evolve alongside tumour progression (32). Specifically, breast cancer tissue consistently exhibits higher stiffness compared to surrounding healthy tissue (32–34), while being accompanied by a concomitant loss of viscoelasticity (32, 34). However, the exact relationship between matrix viscoelasticity and spatial confinement in driving epithelial cell mechanotransduction and migration during cancer progression remains unclear.

To tackle this challenge, we formulated a series of four polyacrylamide (PAAm) hydrogels with independently adjustable Young’s modulus (*E*) and varying degrees of viscous dissipation, spanning the whole spectrum of mechanical properties observed during breast tumour progression (33–35). Termed “soft” (*E* ∼ 0.3 kPa) and “stiff” (*E* ∼3 kPa), these hydrogels were classified as “elastic” (E) or “viscoelastic” (V) based on their stress relaxation profiles and loss tangent (tan(δ)) values, which were either lower or higher than 0.1, respectively (10). Additionally, we employed soft micropatterning techniques to create 1D fibronectin (FN) lines on these hydrogels, facilitating the exploration of the interplay between spatial confinement and matrix viscoelasticity in a microenvironment mimicking certain aspects of the 3D ECM (22, 36). Our results shed light on the intricate interplay between elasticity, viscoelasticity and spatial confinement in orchestrating epithelial cell migration, offering insights into the role of time-dependent matrix mechanics in the context of breast cancer progression.

## Results and Discussion

### Micropatterned viscoelastic polyacrylamide hydrogels to capture the pathophysiology of the breast tissue microenvironment

We engineered new polyacrylamide (PAAm) hydrogels to emulate the viscoelastic properties characteristic of both healthy and cancerous tissue. Notably, PAAm hydrogels serve as a well-established model in mechanobiology research owing to their adaptable and customizable physicochemical properties (37, 38). Differences in elasticity have been documented between normal and tumorous breast human tissues, with values of *E* ≤ 0.5 kPa for non-tumour tissue and *E* ≥ 3 kPa for malignant breast tumours (33–35, 39). Therefore, our initial focus was on fabricating standard linearly elastic hydrogels with Young’s modulus approximately of 0.3 kPa and 3 kPa, representing the stiffness of healthy and tumour breast tissues, respectively. This was achieved by maintaining the Acrylamide (AAm) percentage at ∼4% and doubling the percentage of bisacrylamide (Bis) crosslinker from ∼0.05% to ∼0.1 %, as confirmed by quasi-static nanoindentation experiments. We denoted these hydrogels “soft elastic” and “stiff elastic” (Soft E / Stiff E), respectively (**Figs. 1a and c**).

**Figure 1–.**
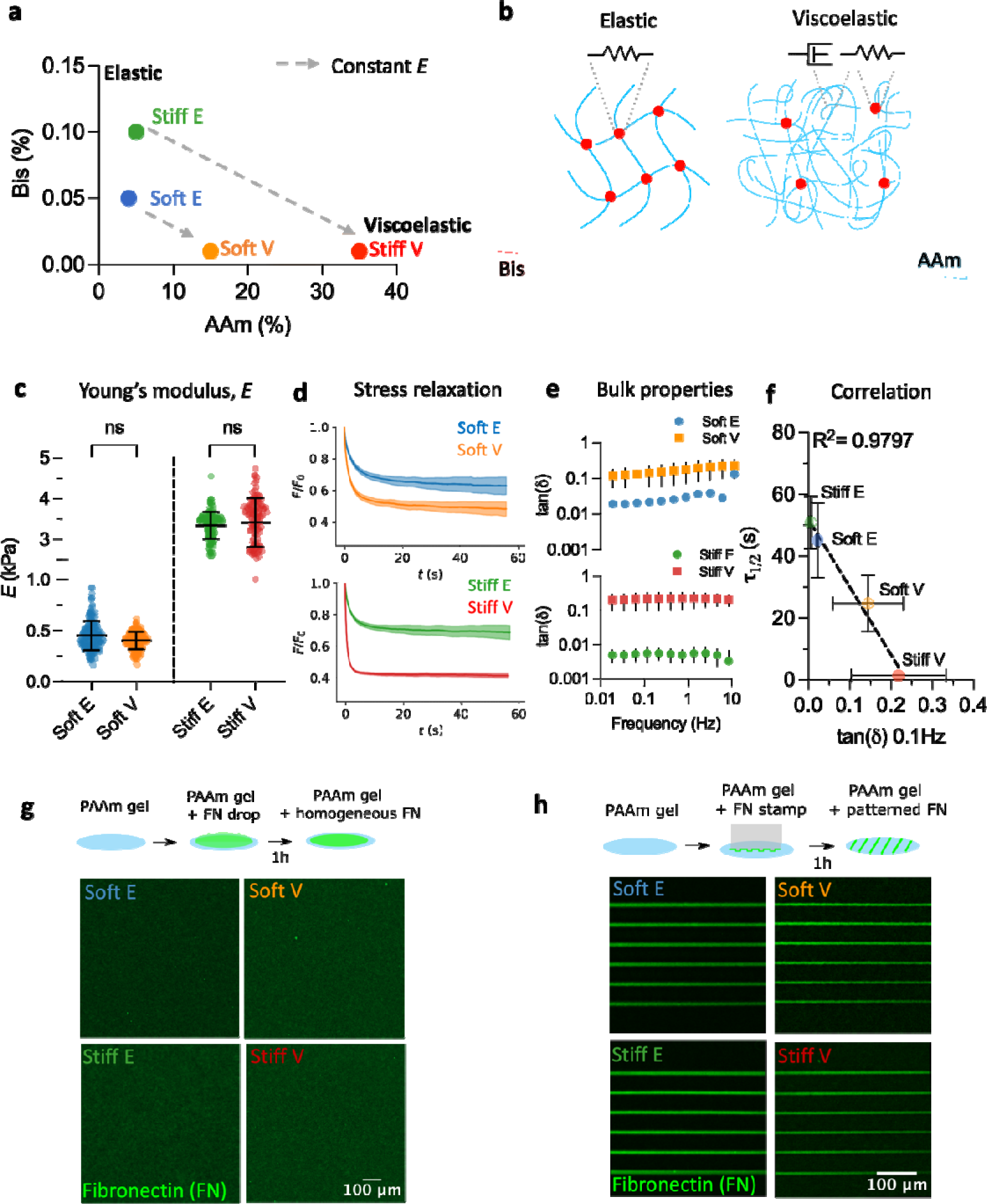
Engineering viscoelastic micropatterned polyacrylamide hydrogels to recapitulate the microenvironments of breast tissue. **(a)** Phase diagram showing the percentages of Bisacrylamide (Bis) crosslinker and Acrylamide (AAm) monomer used to tune the viscoelastic properties of polyacrylamide hydrogels used in this study. Viscoelastic hydrogels have a high AAm to Bis ratio compared to elastic hydrogels. Soft E = soft elastic, Soft V = soft viscoelastic, Stiff E = stiff elastic, Stiff V = stiff viscoelastic. The dashed grey lines connect hydrogels of similar Young’s modulus (*E*) but different viscoelastic properties. **(b)** Schematic representation of the strategy used to obtain elastic and viscoelastic hydrogels with the same initial Young’s modulus. The amount of Bisacrylamide (Bis) is decreased while concurrently increasing the amount of Acrylamide (Aam) to favour physical entanglements. Red dots represent chemical crosslinks, idealised by an elastic spring. Chain entanglements are idealised by a viscous dashpot. **(c)** Young’s modulus (*E*) of hydrogels used in this work. Each point represents a single indentation, with at least 121 indentations (121 ≥ n ≥ 173) from three independently prepared samples. ns p = 0.3427 (Soft group) and p = 0.1453 (Stiff group), two-way ANOVA with Bonferroni’s multiple comparisons test. **(d)** Average stress relaxation profiles of hydrogels used in this work. Curves were obtained by averaging at least 121 individual curves (121 ≥ n ≥ 151) coming from at least two independent experiments. **(e)** tan(δ) obtained from bulk rheology oscillatory sweeps of hydrogels used in this work. Data has been averaged over three independent samples. **(f)** Correlation between the relaxation half-time (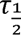) obtained from nanoindentation experiments and the tan(δ) at 0.1 Hz obtained from bulk rheology experiments for the same data shown in d and e (R^2^= 0.9797, mean ± SD). Note that elastic hydrogels dissipated less than 50% of the original stress, so the relaxation half time was taken from the time point resulting in a stress value as close as possible to 50%. **(g)** Representative images of homogeneous fibronectin (FN) coating on elastic and viscoelastic polyacrylamide (PAAm) hydrogels. **(h)** Representative images of micropatterned FN coating on elastic and viscoelastic PAAm hydrogels.

Our next objective was to develop hydrogels with nearly identical Young’s modulus but enhanced viscoelasticity, aiming to dissect the influence of energy dissipation on cellular behaviour. Various strategies have been proposed to independently control the stiffness and viscoelasticity of PAAm hydrogels, including integrating viscous pre-polymerized linear acrylamide into an elastic network (14, 40), adjusting the initiator and activator concentrations (20), or modifying the Aam-to-Bis ratio (16). We recently refined the latter approach to generate isoelastic matrices with diverse dissipative profiles (18), where viscoelasticity is enhanced by increasing the concentration of monomer while concurrently decreasing the concentration of crosslinker (**Fig. 1b**). In this scenario, the network’s elasticity is governed by the monomer concentration, while the low concentration of crosslinker favours physical entanglements between long and loosely bound polyacrylamide chains, providing viscoelastic characteristics to the material (16, 18). By maintaining the Bis percentage at ∼0.01 % and increasing the AAm percentage from 15% to 35%, we produced viscoelastic PAAm hydrogels with initial *E* akin to that of elastic hydrogels (**Figs. 1a and c**). We termed these gels “soft viscoelastic” (Soft V) and “stiff viscoelastic” (Stiff V). We opted for this terminology (i.e., soft vs stiff) for consistency with recent literature (35). However, it is important to note that the hydrogels used in this study are soft in absolute terms (1, 10). Hence, the distinction between “soft” and “stiff” more accurately reflects a tenfold relative change in Young’s modulus between the two hydrogel groups.

To confirm the augmented dissipative properties of the viscoelastic hydrogels, we conducted nanoindentation stress-relaxation experiments, as well as bulk rheology oscillatory analyses. Nanoindentation stress-relaxation experiments, aimed at assessing viscoelasticity at similar force and length scales experienced by cells (41–43), entailed applying a step indentation of ∼10% of the bead’s radius for 60 seconds, resulting in a physiologically relevant 2D strain of approximately 7 % (Materials and Methods) (17). Viscoelastic hydrogels demonstrated enhanced and faster stress relaxation compared to their elastic counterparts (**Fig. 1d**), with viscoelastic hydrogels dissipating more than 50 % of the initial force during the experiment **(Supp. Figs. S1a and b)**. All hydrogels reached a plateau in stress relaxation within 60s, with a fraction of the stress remaining undissipated. Notably, our hydrogels exhibited behaviour characteristic of viscoelastic solids, rather than fluids, across the physiologically-relevant mechanosensitive time scale under investigation, experimentally estimated to be over tens of seconds from traction force fluctuations (14). It is important to note that in an ideal elastic solid, there is no energy dissipation, and therefore the stress remains constant over time under the applied strain. As a result, the stress relaxation half-time is considered infinite in ideal elastic materials. Importantly, the relaxation dynamics and associated time scales exhibit significant variability depending on the measurement scale, spanning from bulk measurements (e.g., through bulk compression tests) to local ones (e.g., via nanoindentation) (11, 41). As a result, the diversity in measurement techniques present considerable obstacles to directly compare relaxation timescales of hydrogels across different studies.

We further confirmed the viscoelastic properties of the bulk hydrogels through oscillatory bulk rheology tests. Specifically, we assessed the ratio between the loss modulus and the storage modulus, commonly referred to as the loss tangent or tan(δ), which serves as an indicator of material viscosity (11). Interestingly, it has been shown that soft viscoelastic ECMs exhibit high tan(δ) values, typically exceeding 0.1 (10, 11). Our analyses revealed significantly higher tan(δ) values for viscoelastic hydrogels compared to their elastic counterparts across the entire frequency range tested (**Fig. 1e**), thus confirming their enhanced dissipative properties.

Correlation analysis between the average time for the stress to reach half of its original value (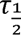) obtained from nanoindentation experiments and the tan(δ) at 0.1 Hz obtained from bulk oscillatory experiments revealed a robust linear relationship between these variables (**Fig. 1f**, R^2^ = 0.9797). This suggests that both metrics serve as reliable indicators of the viscoelastic behaviour of the hydrogels developed in this study. We selected 0.1 Hz as it corresponds to a similar equivalent timescale as the one in stress relaxation experiments (tens of seconds), rendering it a biologically relevant frequency, as previously proposed (14).

We further observed a good correlation between the stress relaxation amplitude (i.e., the percentage of energy dissipated over the experiment’s time) and the tan(δ) at 0.1 Hz, corroborating the enhanced dissipative properties of viscoelastic hydrogels compared to their elastic counterparts **(Supp. Fig. S1c)**.

Given the non-adhesive nature of PAAm hydrogels, functionalisation with ECM proteins is essential to allow for cell adhesion (37, 44). Conventional PAAm gel functionalisation typically involves UV activation of the hetero-bifunctional crosslinker Sulfo-SANPAH (37). However, the compound’s high instability in aqueous solutions often leads to substantial variability in protein crosslinking efficiency on the hydrogel surface (45, 46). This instability poses a significant challenge, particularly for micropatterning hydrogels, where the time required to dry the hydrogel surface exceeds the compound’s half-life (45). To address this issue, we adapted a previously published protocol enabling robust binding of ECM proteins to PAAm hydrogels through the incorporation of reactive aldehyde groups within the network (45, 47). This modification facilitates the covalent conjugation of EMC proteins onto the hydrogel surface regardless of the hydrogel mechanical properties, eliminating the need for additional activation steps post-gelation (45, 47). Fibronectin (FN), an important component of the interstitial breast ECM whose upregulation is associated to breast cancer progression (48), was conjugated onto the hydrogels either homogeneously or through micropatterning narrow lines of approximately 5 µm in width via microcontact printing (Materials and Methods) (**Figs. 1g and h**).

In summary, we have established a system that faithfully replicates the entire spectrum of mechanical properties observed during breast cancer progression, specifically the transition from a soft viscoelastic tissue to a stiff elastic one (32, 34). Importantly, this system enables the independent investigation of the effects of elasticity, viscoelasticity, and spatial confinement on cell behaviour.

### Viscoelasticity modulates cell spreading, focal adhesions and YAP nuclear import in opposite directions on soft and stiff substrates

We first investigated the impact of matrix viscoelasticity on key cellular processes, including cell spreading, adhesion formation and nuclear YAP translocation. Cell response to matrix viscoelasticity exhibits cell-type dependency and heterogeneity (49, 50). To delineate epithelial cell response to matrix energy dissipation, we employed MCF-10A cells, a well-established model for the breast epithelium (51). Focussing on single cells devoid of cell-cell contacts allowed us to isolate ECM mechanics-mediated mechanotransduction effects (52).

After culturing cells on viscoelastic PAAm gels for at least 24h, we observed significant differences in cell spreading area and morphology as a function of substrate viscoelastic properties, as quantified by immunofluorescence (**Figs. 2a-c**). Notably, cell spreading area increased with enhanced viscoelasticity in soft hydrogels and decreased with increased viscoelasticity in stiff hydrogels (**Figs. 2a and b**). Overall, cell spreading area was minimal on stiff V and soft E hydrogels, maximal on stiff E hydrogels, and intermediate on soft V hydrogels. Cell circularity exhibited an opposite trend to cell spreading area (**Figs. 2a and c**), suggesting a reciprocal relationship. To further validate that mammary epithelial cells perceive stiff elastic hydrogels as substrates with high stiffness, we cultured MCF-10A cells on FN-coated glass coverslips. Interestingly, we observed no significant difference in cell spreading area and cell circularity compared to cells cultured on stiff E hydrogels **(Supp. Fig. S2)**, confirming that breast epithelial cells perceive stiff elastic hydrogels as substrates exhibiting infinite rigidity.

**Figure 2–.**
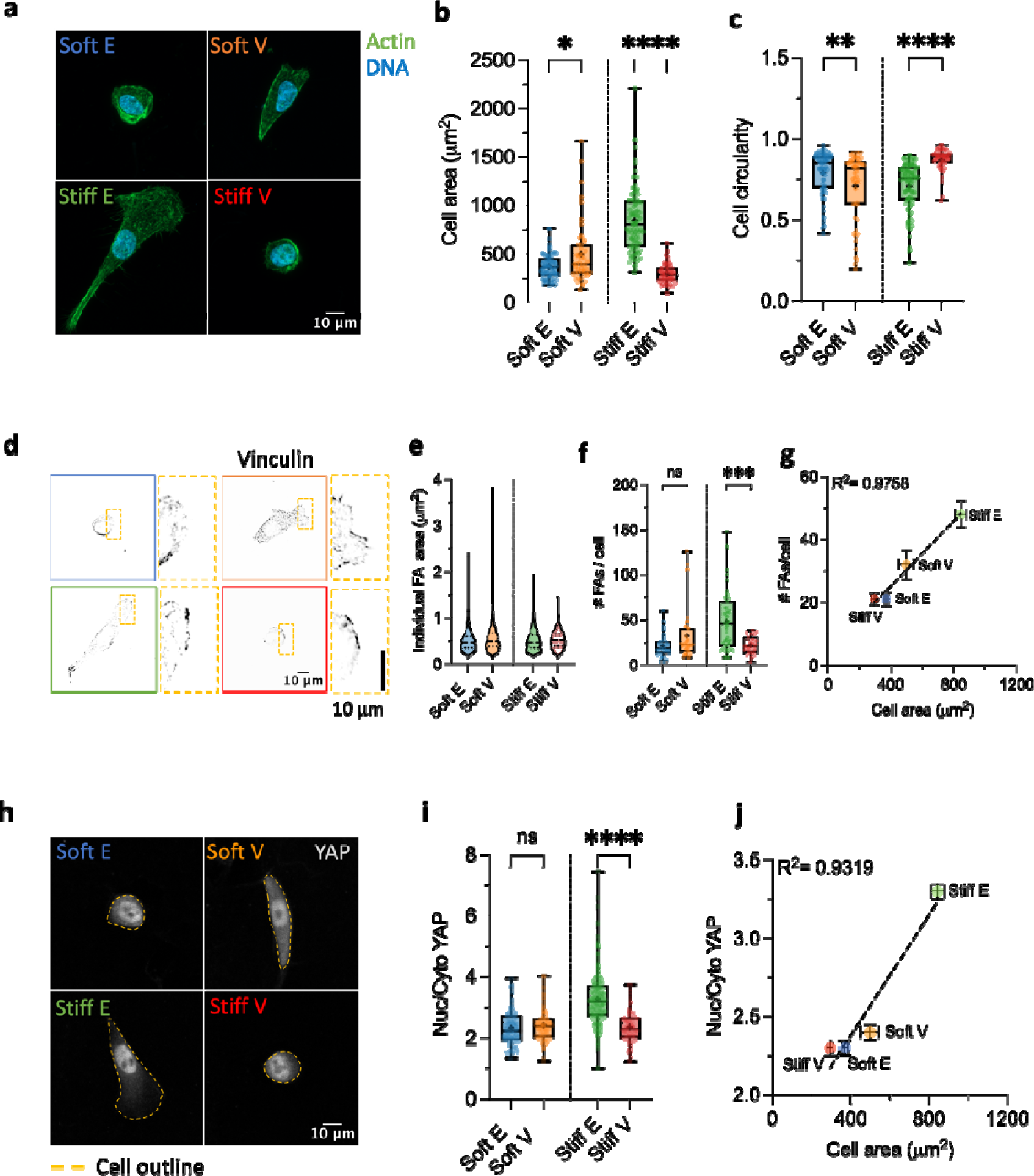
Viscoelasticity modulates cell spreading, focal adhesions and YAP nuclear import in opposite directions on soft and stiff substrates. **(a)** Representative Actin/DNA images of typical MCF-10A cell morphologies observed on elastic and viscoelastic polyacrylamide matrices. **(b)** Quantification of MCF-10A cell spreading area on soft E (n = 74 cells), Soft V (n=51 cells), Stiff E (n=100 cells) and stiff V (n = 49 cells) hydrogels from at least two independent experiments. * p = 0.0148, **** p<0.0001, two-way ANOVA with Bonferroni’s multiple comparisons test. **(c)** Quantification of MCF-10A cell circularity on soft E (n = 74 cells), Soft V (n=51 cells), Stiff E (n=100 cells) and stiff V (n = 49 cells) hydrogels from at least two independent experiments. **p = 0.0061, **** p<0.0001, two-way ANOVA with Bonferroni’s multiple comparisons test. **(d)** Representative focal adhesions (Vinculin) images of MCF-10A cells cultured on elastic and viscoelastic polyacrylamide hydrogels. **(e)** Distribution of individual focal adhesion area of MCF-10A cells cultured on Soft E (n = 685 adhesions), Soft V (n=1301 adhesions), Stiff E (n=3055 adhesions) and Stiff V (n=580 adhesions) hydrogels. Data was obtained from at least two independent experiments. **(f)** Quantification of the number of focal adhesions per cell (#FAs/cell) of MCF-10A cells cultured on soft E (n = 33 cells), soft V (n=41 cells), Stiff E (n=58 cells) and stiff V (n=27 cells) hydrogels from at least two independent experiments. ns p = 0.1413, ****p<0.0001, two-way ANOVA with Bonferroni’s multiple comparisons test. **(g)** Plotting the #FAs/cell vs the cell spreading area reveals a linear relationship between the two variables (R^2^ = 0.9758, data shown as mean ±SEM). **(h)** Representative YAP images of MCF-10A cells cultured on elastic and viscoelastic polyacrylamide hydrogels. The cell’s outline is highlighted by a dashed yellow line. Note the absence of almost any cytoplasmic YAP on stiff E matrices compared to the other conditions. **(i)** Quantification of the Nuclear to Cytoplasmic (Nuc/Cyto) YAP ratio of MCF-10A cells cultured on soft E (n=157 cells), Soft V (n=128 cells), Stiff E (n=320 cells), Stiff V (n=113 cells) hydrogels from at least two independent experiments. ns p = 0.5161, ****p<0.0001, two-way ANOVA with Bonferroni’s multiple comparisons test. **(j)** Nuc/Cyto YAP ratio increases linearly with cell spreading area on viscoelastic polyacrylamide hydrogels (R^2^ = 0.9319, data shown as mean ±SEM).

While increased matrix rigidity traditionally promotes cell spreading (4), conflicting results have been reported regarding the inhibition or enhancement of cell spreading ability with increased viscoelasticity (13–15, 49, 53, 54). Our findings provide insight into these results by demonstrating that increased viscoelasticity promotes cell spreading when the initial substrate stiffness is low (∼*E* < 1 kPa) and inhibits it when the initial substrate stiffness is high (∼*E* > 1 kPa) (13, 53). Interestingly, our results are consistent with theoretical predictions using molecular clutches interacting with a viscoelastic substrate, which suggest a potential regime at low stiffness and high ligand density in which cell spreading is enhanced (13, 53). To confirm the role of ligand density within our experimental setup, we reduced the FN concentration utilized for functionalizing PAAm hydrogels by sevenfold, from 70 μg/mL to 10 μg/mL. This adjustment abrogated the previously observed increase in cell spreading on soft V matrices compared to soft E ones, while still preserving the diminished spreading observed on Stiff V matrices in contrast to stiff E matrices **(Supp. Fig. S3)**, consistent with theoretical simulations (13, 53).

Cell spreading in 2D serves as a proxy for mechanically-driven increased cytoskeletal tension and cell contractility (55). Focusing on vinculin, a focal adhesion (FA) protein within the molecular clutch axis (53, 56, 57), we found no biologically meaningful variations of individual FA area across the different matrices (**Figs. 2d and e**), suggesting that FA reinforcement does not occur over the range of ECM viscoelastic properties explored (58). However, our findings revealed that the number of FAs per cell increased linearly with the projected cell spreading area (**Figs. 2f and g**, R^2^=0.9758). These results demonstrate that increased matrix viscoelasticity enhances cell contractility at low initial stiffness, while conversely diminishing cell contractility when the initial stiffness of the ECM is elevated (55). Meanwhile, FA density remained almost constant, suggesting that elevated matrix viscoelasticity supports cell adhesion.

On 2D ECMs, the mechanical tension of actin stress fibres acts on the nucleus through the Linker of Nucleoskeleton and Cytoskeleton (LINC) complex, facilitating YAP nuclear import —a fundamental transcriptional regulator mediating mechanotransduction. Interestingly, we observed predominantly nuclear YAP in all conditions, with Nuclear (Nuc)/Cytoplasmic (Cyto) ratios exceeding 2 (**Figs. 2h and i**). We hypothesised that the heightened FN concentration utilized in this work likely contributed to the elevated basal level of nuclear YAP across all conditions, a phenomenon recognized to enhance nuclear YAP irrespective of ECM mechanical properties (59). To confirm this, we cultured cells on the same viscoelastic hydrogels but employed a sevenfold lower FN concentration, from 70 μg/mL to 10 μg/mL, resulting in the abolishment of YAP nuclear localization across all conditions except for stiff E matrices, as anticipated **(Supp. Fig. S4)**. Nevertheless, nuclear YAP import remained responsive to ECM viscoelastic properties even at high FN ligand density, exhibiting a linear increase with cell projected area (**Fig. 2j**, R^2^=0.9319) and with the number of FAs per cell **(Supp. Fig. S5)**. These results underscore that cellular contractility governs nuclear YAP import in response to matrix viscoelastic properties.

Altogether, our findings underscore the role of matrix viscoelasticity, in addition to stiffness, as a key mediator of mechanotransduction in ECMs mimicking both physiological and pathological conditions in mammary tissues.

### Viscoelasticity enhances migration speed and persistence on soft substrate, while impeding them on stiff substrates via actin retrograde flow regulation

Acknowledging the pivotal role of focal adhesions as critical mediators of cell migration through molecular clutch mechanisms (60), we next sought to investigate the influence of ECM viscoelasticity on cell migration. Recent studies have indicated that matrix stress relaxation enhances MCF-10A migration on relatively soft fast-relaxing alginate-reconstituted basement membrane (rBM) hydrogels (*E* ∼2 kPa) compared to their slow-relaxing counterparts (19). While these findings are significant for understanding viscoelastic matrices resembling cancerous breast tissues, questions persist regarding how heightened energy dissipation impacts epithelial cell migration within mechanical microenvironments resembling the stiffness of healthy (*E* ∼0.3 kPa) and cancerous (*E* ∼3 kPa) breast tissue (33–35). Furthermore, recent demonstrations of the evolution of tissue viscoelastic properties during breast tumour progression (32–34) has fuelled our interest in elucidating the role of matrix viscoelasticity in driving breast epithelial cell migration on soft and stiff elastic and viscoelastic ECMs.

We conducted 15-hour time lapse experiments to study the migratory behaviour of MCF-10A cells on elastic and viscoelastic PAAm hydrogels. By tracking individual cells, we quantified their average speed and mean square displacement (MSD) (61). Migration tracks in **Fig. 3a** reveal that enhanced energy dissipation has a dual effect on cell migration **(Supp. Movies 1 to 4)**. Specifically, cells on soft V substrates migrated faster and further than cells on their elastic counterparts (i.e., Soft E), exhibiting both increased average speed and MSD (**Figs. 3b and c**). Conversely, both speed and MSD significantly decreased for cells on stiff V matrices compared to cells on stiff E ones (**Figs. 3d and e**). The slope of the MSD-lag time data yielded the diffusion exponent, α, indicating migration persistence (61). Cells on soft V substrates displayed higher persistence than those on soft E substrates, while cells on stiff E substrates were more persistent than cells on stiff V ones. Additionally, we observed no significant difference in migration speed between cells cultured on soft V substrates compared to those cultured on FN-coated glass coverslips **(Supp. Fig. S6)**, confirming that enhanced viscoelasticity on soft substrates promotes cell migration to similar levels observed on infinitely stiff elastic substrates.

**Figure 3–.**
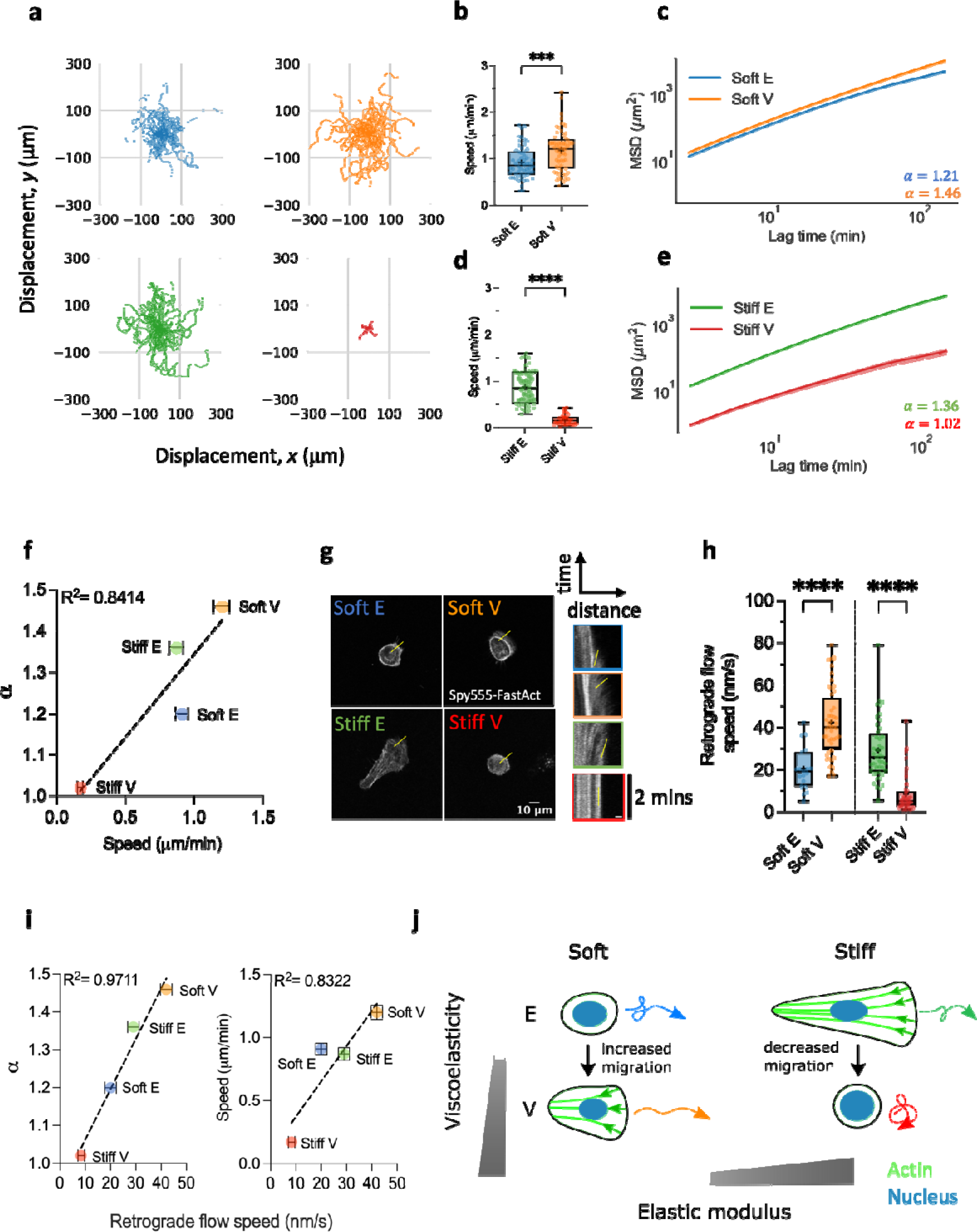
Viscoelasticity enhances migration speed and persistence on soft substrates, while impeding them on stiff substrates via actin retrograde flow regulation. **(a)** Representative *x-y* trajectories of MCF-10A cells on elastic and viscoelastic polyacrylamide hydrogels over 5 hours (n = 48 trajectories for soft E, n = 43 trajectories for Soft V, n = 49 trajectories for Stiff E, n = 34 trajectories for Stiff V). **(b)** MCF-10A cell migration speed on soft elastic (Soft E, n=56 cells) and viscoelastic (Soft V, n = 61 cells) matrices obtained from three independent experiments. *** p = 0.0007, unpaired two-tailed t-test. **(c)** Average mean square displacement (MSD) vs lag time for MCF-10A cells on soft elastic (Soft E) and viscoelastic (Soft V) matrices. The diffusion exponent, α, is shown in the graph. Data is shown as mean ±SEM (n = 48 cells for Soft E, n = 43 cells for Soft V) from three independent experiments. **(d)** MCF-10A cell migration speed on stiff elastic (Stiff E, n=55 cells) and viscoelastic (Stiff V, n = 35 cells) matrices obtained from at least two independent experiments. *** p = 0.0007, unpaired two-tailed t-test. **** p < 0.0001, unpaired two-tailed t-test. **(e)** Average mean square displacement (MSD) vs lag time for cells on stiff elastic (Stiff E) and viscoelastic (Stiff V) matrices. The diffusion exponent, α, is shown in the graph. Data is shown as mean ±SEM (n = 49 cells for Stiff E, n = 34 cells for Stiff V) from at least two independent experiments. **(f)** Diffusion exponent, α, plotted against average cell migration speed. Data is shown as mean ±SEM (R^2^ = 0.8414) for the same number of cells and independent experiments as in b-d (for cell migration speed) and c-e (for MSD). **(g)** Representative images of MCF-10A tagged with live Spy555-FastAct on elastic and viscoelastic polyacrylamide hydrogels. Yellow line shows location where kymographs were computed, on average. Insets show representative kymographs for each condition, with yellow line indicating the slope from which the actin retrograde flow speed is computed. The spatial scale bar in the inset is 2 μm, whereas the temporal scale bar is 2 minutes. **(h)** Quantification of actin retrograde flow speed for MCF-10A cells cultured on Soft E (n = 19 kymographs), Soft V (n=48 kymographs), Stiff E (n=44 kymographs) and Stiff V (n=52 kymographs) hydrogels from at least two independent experiments. **** p < 0.0001, two-way ANOVA with Bonferroni’s multiple comparisons test. **(i)** Diffusion exponent, α, vs actin retrograde flow speed (left, R^2^ = 0.9711); and migration speed vs actin retrograde flow speed (right, R^2^ =0.8322). Data is shown as mean ±SEM for the same number of cells/kymographs as in c-e and h, respectively. **(j)** Schematic summary of experimental findings on how matrix viscoelasticity modulates MCF-10A cell migration speed and persistence via actin retrograde flow speed. Green arrows inside the cells represent actin retrograde flow, whereas colourful arrows outside the cell depict schematic migration trajectories.

Overall, our results reveal that enhanced ECM viscoelasticity can either promote or hinder 2D cell migration depending on initial ECM stiffness. Different cell types have varying number of active molecular motors and clutches that establishes a stiffness optimum for cell migration (7). Consequently, matrix viscoelasticity is likely to affect cell migration in non-trivial ways, depending on the initial stiffness of the substrate, relaxation time scale and ECM ligand type and density (50).

Furthermore, we observed a strong linear correlation between cell persistence and cell speed (**Fig. 3f**, R^2^= 0.8414), a universal characteristic of migrating cells in vitro and in vivo regardless of matrix dimensionality (i.e., 2D vs 3D) and migration mode (62). This relationship is governed by actin polymerization rate, indicating that actin flow speed links the observed universal coupling between cell speed and persistence (62). To delve into the mechanism, we measured actin flow speed at the lamellipodium edge of migrating cells. Cells migrating on stiff E matrices displayed a fan-shaped phenotype with an extended lamellipodium, while cells on soft V matrices exhibited a more compacted morphology with a short lamellipodium. Cells on soft E hydrogels did not develop a prominent lamellipodium, similar to those on stiff V hydrogels **(Supp. Fig. S7** and **Supp. Movies 5 to 8)**. MCF-10A cells were tagged to label fast-polymerizing actin filaments, and cells were imaged at high spatial and temporal resolution for 2 minutes at 1-second intervals (**Fig. 3g** and **Supp. Videos 9 to 12)**. Using kymographs, we quantified the actin retrograde flow speed at the lamellipodium edge for cells on soft V and stiff E matrices, and at random anterior locations for cells on Soft E and stiff V matrices (**Fig. 3h**). Our findings indicated that the retrograde flow speed increased two-fold by adding dissipative components in soft matrices, while in contrary it was impeded by incorporating viscous dissipation in stiff matrices (**Fig. 3h**). Interestingly, we observed a robust linear correlation between the diffusion exponent and the actin retrograde flow speed (**Fig. 3i**, R^2^=0.9711), as well as the migration speed and the actin retrograde flow speed (**Fig. 3i**, R^2^=0.8322), indicating that the viscoelastic properties of the cell substrate regulate cell persistence and speed through modulation of the retrograde flow of actin filaments (62). These results suggest that optimal retrograde flow occurs on soft viscoelastic substrates (*E* ∼0.3 kPa) mimicking the mechanical properties of healthy tissues, when the substrate relaxation time likely aligns with the timescale for clutch binding and its characteristic binding lifetime (53). Conversely, on stiff viscoelastic substrates (*E* ∼3 kPa), viscosity significantly reduces cell spreading (**Fig. 2b**), cell speed (**Fig. 3d**), cell persistence (**Fig. 3e**) and retrograde flow speed (**Fig. 3h**) by impeding the backward movement of actin filaments.

Taken together, our results underscore the influence of substrate viscoelasticity on epithelial cell migration, revealing distinct responses dependent on substrate stiffness. Notably, we observed that shorter stress relaxation times enhance cell migration on soft substrates (*E* ∼0.3 kPa) while significantly impeding it on stiff substrates (*E* ∼3 kPa). At a first sight, our findings may seem to contradict recent results obtained on 2 kPa viscoelastic alginate-rBM hydrogels (19). However, upon comparing cell migration on 2kPa slow-relaxing alginate-rBM hydrogels to migration on elastic PAAm hydrogels of the same elastic modulus, the authors found that both cell speed and directional persistence were higher on purely elastic PAAm hydrogels compared to slow-relaxing alginate-rBM ones (19). Since the viscoelastic hydrogels developed in our work are chemically crosslinked, their bulk relaxation half time is expected to be within the range of several hundred to a few thousand of seconds (11), similarly to slow-relaxing hydrogels reported previously (19). Therefore, our findings align with recent results on the effects of substrate stress relaxation on cell migration on matrices with elastic moduli of 2-3 kPa.

However, our findings also reveal an intriguing regime on softer substrates (*E* ∼0.3 kPa), wherein mammary epithelial cells exhibit optimal migration on soft viscoelastic substrates, recapitulating the native stiffness and stress-relaxing properties of healthy breast tissue. Drawing upon the evolving understanding of tissue viscoelastic properties during breast tumour progression (32–34), these findings underscore the pivotal role of substrate viscoelasticity as a fundamental ECM physical property, profoundly influencing the regulation of cell migration across different stiffness regimes.

### Spatial confinement restricts cell migration on soft matrices regardless of matrix viscoelasticity and promotes migration on stiff matrices in a viscoelasticity-dependent manner

In addition to changes in viscoelasticity within their microenvironment, epithelial mammary cells encounter spatial constraints imposed by neighbouring cells or the ECM (23). Confined spaces have been shown to significantly influence cell migration (24–26, 29, 30, 64), with spatial confinement playing a crucial role in metastatic spreading during cancer progression (23). Interestingly, 1D confinement has been demonstrated to induce a spindle-like morphology and migration phenotype resembling that of cells in soft 3D matrices, effectively capturing certain aspects of 3D cell migration (22). While few previous studies have explored the interplay between substrate stiffness and confinement in terms of cell migration (24, 29, 30), the impact of the coupling between increased matrix viscoelasticity and spatial confinement on cell migration remains elusive.

To address this gap, we micropatterned 5 μm FN lines on all substrates (**Fig. 1h**), corresponding to the upper threshold above which cells lose their uniaxial 1D spindle-like morphology (22). We first studied whether 1D confinement and viscoelasticity could affect the morphology of breast epithelial cells in terms of projected spreading area and aspect ratio. Regardless of matrix viscoelastic properties, our findings show that cells on 5 µm lines adopted a 1D spindle-like morphology with a high aspect ratio (**Figs. 4a-c**) compared to 2D conditions (**Fig. 2a**). However, 1D confinement affected projected cell spreading area differentially depending on ECM viscoelasticity. Specifically, on soft substrates, cell spreading area was not affected by confinement (**Fig. 4d**). Conversely, cell spreading area was significantly reduced on 1D stiff E matrices compared to their 2D counterparts, consistent with results reported on stiff elastic substrates (22, 65) (**Fig. 4e**), indicating that cells are restricted when confined to 1D lines on stiff E matrices. Similarly to soft substrates, cell spreading area was not affected on stiff V matrices as compared to their 2D counterpart. However, cells became elongated and spread along the pattern compared to their circular unpolarised morphology on 2D matrices (**Fig. 4e**).

**Figure 4–.**
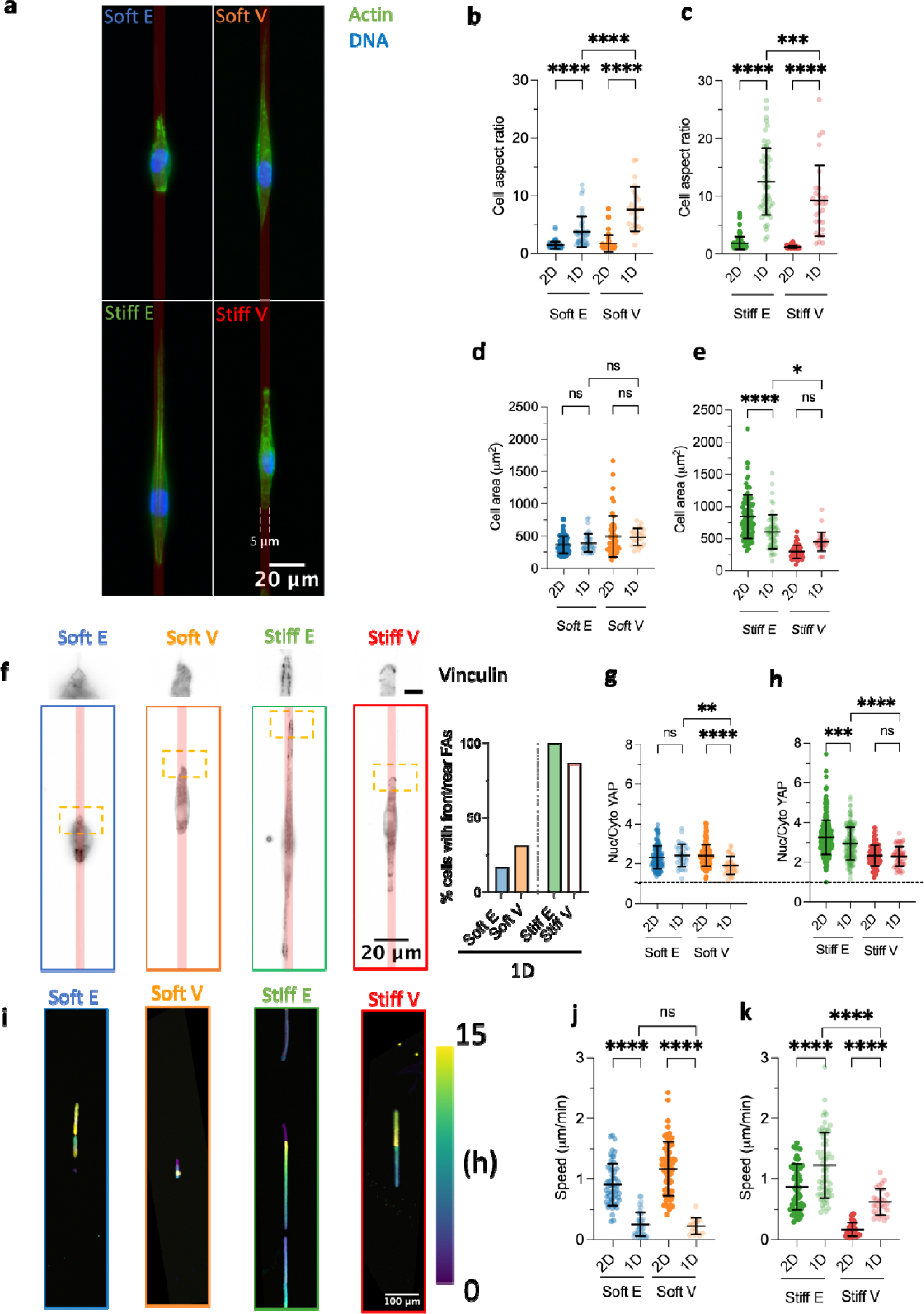
Spatial confinement restricts cell migration on soft matrices regardless of viscoelasticity and promotes migration on stiff matrices in a viscoelasticity-dependent fashion. **(a)** Representative Actin/DNA images of confined MCF-10A cells on 5 μm fibronectin lines on elastic and viscoelastic polyacrylamide hydrogels. Fibronectin lines are schematically represented in red for clarity. **(b)** Quantification of MCF-10A cell aspect ratio on 2D and 1D soft elastic (Soft E) and viscoelastic (Soft V) matrices (n = 74 cells for Soft E 2D, n = 40 cells for soft E 1D, n = 50 cells for Soft V 2D, n = 24 cells for soft V 1D) from at least two independent experiments. **** p < 0.0001, two-way ANOVA with Tukey’s multiple comparisons test. **(c)** Quantification of MCF-10A cell aspect ratio on 2D and 1D stiff elastic (Stiff E) and viscoelastic (Stiff V) matrices (n = 100 cells for Stiff E 2D, n = 59 cells for Stiff E 1D, n = 48 cells for Stiff V 2D, n = 29 cells for Stiff V 1D) from at least two independent experiments. ****p < 0.0001, ***p = 0.0006, two-way ANOVA with Tukey’s multiple comparisons test. **(d)** Quantification of MCF-10A cell spreading area on 2D and 1D soft elastic (Soft E) and viscoelastic (Soft V) matrices (n = 74 cells for Soft E 2D, n = 40 cells for soft E 1D, n = 50 cells for Soft V 2D, n = 24 cells for soft V 1D) from at least two independent experiments. ns p > 0.05, two-way ANOVA with Tukey’s multiple comparisons test. **(e)** Quantification of MCF-10A cell spreading area on 2D and 1D stiff elastic (Stiff E) and viscoelastic (Stiff V) matrices (n = 100 cells for Stiff E 2D, n = 59 cells for Stiff E 1D, n = 48 cells for Stiff V 2D, n = 29 cells for Stiff V 1D) from at least two independent experiments. ****p<0.0001, *p=0.0473, ns p =0.0588, two-way ANOVA with Tukey’s multiple comparisons test. **(f)** Representative focal adhesions (Vinculin) images of confined MCF-10A cells on 5 μm fibronectin lines on elastic and viscoelastic polyacrylamide hydrogels. Fibronectin lines are schematically represented in red for clarity. Scale bar in the inset is 5 μm. Bar graph shows the percentage of cells forming front and rear vinculin adhesions pooled from two independent experiments. **(g)** Quantification of nuclear to cytoplasmic (Nuc/Cyto) YAP ratio on 2D and 1D soft elastic (Soft E) and viscoelastic (Soft V) matrices (n = 157 cells for Soft E 2D, n = 35 cells for soft E 1D, n = 128 cells for Soft V 2D, n = 36 cells for soft V 1D) from at least two independent experiments. Dashed line indicates a Nuc/Cyto YAP ratio of 1. ****p < 0.0001, **p = 0.001, two-way ANOVA with Tukey’s multiple comparisons test. **(h)** Quantification of nuclear to cytoplasmic (Nuc/Cyto) YAP ratio on 2D and 1D stiff elastic (Stiff E) and viscoelastic (Stiff V) matrices (n=320 cells for Stiff E 2D, n = 150 cells for stiff E 1D, n=113 cells for stiff V 2D, n = 56 cells for stiff V 1D) from at least two independent experiments. ****p<0.0001, ***p=0.0003, ns p = 0.9911, two-way ANOVA with Tukey’s multiple comparisons test. **(i)** Representative temporal colour coded time lapses of MCF-10A migrating on 5 μm fibronectin lines. **(j)** Quantification of MCF-10A cell migration speed on 2D and 1D soft elastic (Soft E) and viscoelastic (Soft V) matrices (n=56 cells for soft E 2D, n = 38 cells for Soft E 1D, n = 61 cells for Soft V 2D, n = 14 cells for soft V 1D) from at least two independent experiments. ****p < 0.0001, ns p = 0.9911, two-way ANOVA with Tukey’s multiple comparisons test. **(k)** Quantification of MCF-10A cell migration speed on 2D and 1D stiff elastic (Stiff E) and viscoelastic (Stiff V) matrices (n = 55 cells for Stiff E 2D, n = 61 cells for Stiff E 1D, n = 35 cells for Stiff V 2D, n = 26 cells for Stiff V 1D) from at least two independent experiments. ****p <0.0001, two-way ANOVA with Tukey’s multiple comparisons test.

When comparing cell aspect ratio under confined 1D conditions, we observed that cells exhibited greater elongation on soft V hydrogels compared to their elastic counterparts (i.e., soft E) (**Fig. 4b**). Conversely, cells displayed increased elongation on stiff E conditions compared to their viscoelastic (i.e., Stiff V) counterparts. This suggests that breast epithelial cells adapt their morphology in response to the viscoelastic properties of the matrix even under confinement. Additionally, when assessing cell spreading area within 1D confined conditions, we found no significant differences between soft E and soft V matrices (**Fig. 4d**). However, cells on 1D stiff E matrices remained larger than those on 1D stiff V matrices, though to a lesser extent than observed in 2D conditions (**Fig 4e**). This indicates that 1D confinement leads cells to exhibit similar spreading area regardless of the underlying matrix

We further investigated how spatial confinement affects focal adhesions on viscoelastic ECMs, focussing on the spatial localisation of FAs. The formation and disengagement of front-rear FAs have been associated with the ability of cells to break symmetry and migrate when confined on 1D lines on stiff elastic PAAm matrices via a stick-slip mechanism (36). Regardless of matrix energy dissipation, cells on soft matrices rarely formed front-rear FAs, whereas most cells on stiff matrices were able to form front-rear adhesions (**Fig. 4f**).

Interestingly, changes in morphology and adhesions did not strongly change nuclear YAP localisation, which either remained constant or slightly decreased as compared to 2D unconfined ECMs (**Figure 4g and h and Supp. Fig. S8)**.

We then investigated how the observed morphological changes and adhesion phenotype would affect 1D cell migration on viscoelastic ECMs. Migration along 1D lines is an accepted reductionist model to mimic in vivo migration conditions involving confined spaces, such as within dense tissues, along blood vessels, ECM fibres or muscle fibres (23). However, 1D lines are usually produced on stiff glass coverslips (65) or at most on elastic PAAm hydrogels with stiffness in the range of tens of kPa (36), leaving unexplored how matrix-mimetic viscoelastic properties affect 1D confined cell migration on soft substrates. We therefore followed cell migration using time lapse microscopy for 15 hours and observed that migration was abrogated for cells on soft 1D ECMs compared to their 2D counterparts, regardless of viscoelasticity, as indicated by a significant drop in speed (**Figs. 4i and j and Supp. Movies 13-14)**. Conversely, cell migration speed was significantly increased on 1D stiff E and stiff V matrices compared to 2D unconfined conditions (**Figs. 4i and k and Supp. Movies 15-16).**

Previous studies have reported that confinement can either enhance (22) or hinder (25, 65) cell migration on stiff substrates, depending on the cell type. Furthermore, it has been demonstrated that the relationship between cell speed and confinement is influenced by ECM stiffness (24, 29). Our findings agree with previous results, where a decrease in migration was observed in soft and narrow collagen microchannels compared to wider channels of similar mechanical properties (29), and conversely, an increase in migration speed was observed in stiff and narrow collagen microchannels compared to wider channels of similar mechanical properties (29).

Taken together, our results demonstrate that 1D spatial confinement elicits opposite effects on cell migration depending on the stiffness of the matrices. Interestingly, we observed that viscoelasticity does not mediate migration speed in soft 1D ECMs (**Fig. 4j**), suggesting that confinement may mask variations in mechanical properties when the stiffness of the matrix is low. However, viscoelasticity does modulate confined migration in stiff matrices, with cells exhibiting optimal migration in stiff E matrices resembling the tumour microenvironment, and slower migration rates in stiff V matrices (**Fig. 4k**).

The inability of epithelial cells on soft 1D matrices to form front rear FAs implies a deficiency in breaking their own symmetry and polarize, thus impairing their migration. In contrast, cells on stiff substrates can stochastically break their symmetry by disassembling front-rear FAs, initiating migration. These findings are consistent with a one-dimensional (1D) motor clutch-based model, emphasizing the importance of asymmetry in the number of bound clutches between both cell extremities (36). At the cell end with fewer bound molecular clutches, a higher traction force per clutch is sustained, resulting in catastrophic rupture of the cell-substrate adhesions on this side and a subsequent translocation of the cell towards the opposite side. We anticipate that integrating ECM viscoelastic properties across various stiffness regimes into models describing 1D cell migration will offer insights into how migration mechanisms transition from 2D unconfined to 1D confined conditions.

## Conclusion and outlook

This study presents a facile and adaptable method for manipulating viscoelasticity, elasticity, and spatial confinement within polyacrylamide hydrogels. While previous works on viscoelastic matrices (13, 14, 19) have predominantly focused on elasticity ranges above 2-3 kPa, we designed viscoelastic hydrogels with remarkably low elasticities (0.3 ≤ *E* ≤ 3 kPa) to accurately mimic specific aspects of cancer progression, such as the transition from a soft viscoelastic matrix to a stiff elastic matrix.

Mechanistically, our findings revealed that viscoelasticity exerts contrasting effects on cell spreading, focal adhesions, and YAP nuclear import depending on substrate stiffness. Notably, nuclear YAP import remains highly responsive to ECM viscoelastic properties, displaying a linear correlation with cell projected area. These findings underscore the pivotal role of cell contractility in mediating nuclear YAP import in response to matrix viscoelastic properties, complementing the well-established essential involvement of YAP in various aspects of breast cancer development (66). While previous studies have emphasized the role of matrix stiffness in the regulation of metastatic breast cancer cells through YAP signalling (67), our study highlights the pivotal role of substrate viscoelasticity in regulating cell mechanotransduction and migration. Specifically, we demonstrate that viscoelasticity enhances migration speed and persistence on soft substrate, while impeding these processes on stiff substrates via actin retrograde flow regulation.

Although these results were obtained in two dimensions, the combination of these viscoelastic hydrogels with protein micropatterning provides a platform to further investigate the interplay between spatial confinement and viscoelasticity and to capture certain aspects of 3D cell migration. Our findings indicate that spatial confinement restricts cell migration on soft matrices regardless of viscoelasticity and promotes migration on stiff matrices in a viscoelasticity-dependent manner. Altogether, these findings suggest a complex interplay between viscoelastic properties and dimensionality of the surrounding matrix in the regulation of cell migration. Our results derived from 2D experiments with very low elasticities are consistent with a motor clutch-based model substrate wherein viscoelasticity regulates the lifetimes and the number of engaged clutches (19). However, they also indicate that heightened levels of spatial confinement obscure the viscoelastic properties of the substrate, particularly at extremely low stiffness values, by reducing both the number and distribution of cell-substrate interactions, which constitute a critical aspect of the molecular clutch model.

These findings highlight the significance of incorporating stress relaxation as a critical parameter when studying complex cellular behaviours and mechanotransduction signals within human tissues. This approach provides a platform to further decouple the elastic modulus from matrix viscoelastic properties and the level of imposed spatial confinement, that may be useful in a variety of applications involving studying cell migration in tissue-mimetic viscoelastic hydrogels.

## Materials and Methods

### Coverslips silanization

The silanization procedure for coverslips followed previously established protocols (38, 68). Briefly, 22 mm coverslips (VWR) were incubated with 0.1 M NaOH for 5 minutes, followed by three rinses in Milli-Q water, the last time for 5 minutes. Subsequently, coverslips were dried using a gentle nitrogen flow. Next, 15 µL of 3-(Acryloyloxy)propyltrimethoxysilane (Alfa Aesar) per coverslip were pipetted onto a glass plate, and each coverslip was flipped to place the activated side on the drop for 1 hour. Afterwards, coverslips were then rinsed three times in Milli-Q water, gently dried with nitrogen, and stored at 4°C until further use.

### Fabrication of polyacrylamide (PAAm) hydrogels

All reagents for PAAm hydrogels were purchased from Merck. PAAm hydrogels with adjustable viscoelastic properties were fabricated by mixing varying amounts of 40% Acrylamide, 2% Bisacrylamide and Milli-Q water in a 1mL Eppendorf tube. The solutions were gently vortexed to ensure thorough mixing without introducing oxygen. To introduce reactive aldehyde groups for protein binding, a solution of oxidised N-hydroxyethyl acrylamide (OHEA) was prepared as previously outlined (47) and an equal volume of 10 μL added to each prepolymer solution. Each Eppendorf was again gently vortexed to ensure homogeneous mixing of the OHEA within the prepolymer solutions. To initiate polymerization, the same volumes of 100% tetramethylethylenediamine (TEMED) (2.5 μL), 10% ammonium persulfate (APS) in milli-Q water (7.5μL) were added to each Eppendorf. The solutions were gently vortexed one last time. For the formation of thin hydrogel films immobilized on glass coverslips, 30μL of each prepolymer solution were pipetted on a hydrophobic flexible Polychlorotrifluoroethylene (PCTFE) sheet (Agar Scientific) and covered with a 22mm acrylsilanized glass coverslip. Gelation was allowed to occur for 30 minutes at room temperature, after which the PCTFE-coverslip sandwich was submerged in cold (4 °C) milli-Q water for 1h before peeling the PCTFE sheet off. Hydrogels were then rinsed three times with milli-Q water to remove any unreacted groups and left to swell at 4°C overnight in milli-Q water before protein functionalisation. Hydrogel compositions are summarized in **Supp. Table 1**.

### Homogeneous protein functionalisation

Following swelling, hydrogels underwent a thorough rinsing three times with Milli-Q water. Excess water was gently removed from the hydrogel surface by blotting the side of each coverslip against tissue paper, ensuring that the hydrogel surfaces were not completely dry. Subsequently, 100 µL of 70 µg/mL (high concentration) or 10 µg/mL (low concentration) solution of human fibronectin (Yo Proteins) in Milli-Q water was pipetted onto the hydrogel surfaces and allowed to evaporate under airflow. This process facilitated the covalent binding of the primary amines of the fibronectin with the aldehyde groups present in the PAAm hydrogels (45, 47). The samples were then rehydrated in Milli-Q water for 5 minutes and rinsed three times with Milli-Q water to eliminate unbound protein.

### Microstamp fabrication and protein micropatterning

Microstamps featuring 5 µm lines were fabricated using a silicon master obtained through deep reactive-ion etching from a chromium photomask (Toppan Photomask). The silicon master underwent passivation with a fluorosilane (tridecafluoro-1,1,2,2-tetrahydrooctyl-1-trichlorosilane; Gelest) for 30 minutes in a desiccator. Subsequently, polydimethylsiloxane (PDMS) (Sylgard 184 Silicone Elastomer Kit; Dow Corning) mixed at 10:1 w:w ratio (base:crosslinker) was poured onto the master to achieve a height of approximately 1cm and degassed to eliminate any bubbles. The PDMS was cured for a minimum of 4 hours at 60°C. Following this, the PDMS layer was carefully detached from the master, and stamps cut into approximately 1 cm^3^ cubes. PAAm hydrogels were microprinted based on a previously published protocol (38, 68). Briefly, stamps were sonicated in a solution of 5% decon 90 (decon) in milli-Q water for 15 minutes, thoroughly rinsed under water, and sonicated again in a solution of 70% isopropanol in Milli-Q water for 15 minutes. The stamps were then gently dried under nitrogen flow. PDMS stamps were activated using an ultraviolet/O_3_ oven (UV-Ozone photoreactor PR-100, UVP Products) for 8 minutes and incubated with a solution of 70μg/mL fibronectin in Milli-Q water for 1 hour at room temperature. Subsequently, PDMS stamps were gently dried under nitrogen and placed in contact with dried hydrogel surfaces for 1 hour at room temperature. To facilitate stamp detachment from hydrogels, hydrogels/stamps composites were submerged in Milli-Q water for 10 minutes, and stamps carefully removed. To visualise the integrity of patterns during protocol optimisation using fluorescence microscopy, 35 µg/mL of Alexa Fluor 488 or 647 fibrinogen (ThermoFisher) were added to the fibronectin solution.

### Determination of the Young’s modulus

Nanoindentation measurements were conducted using a fiber-optic based nanoindentation device (Chiaro, Optics 11 Life) mounted on top of an inverted optical microscope (Axiovert 200M, Zeiss), following a standardised protocol (43). The hydrogel Young’s modulus (*E*) was determined by moving the probe at a constant speed of 2 µm/s over a vertical range of 10 µm (displacement control). For soft hydrogels, a cantilever with stiffness (*k*) of 0.028 N/m equipped with a spherical bead of 26 µm in radius (*R*) was used. In the case of stiff hydrogels, a cantilever with a *k* of 0.52 N/m and R of 27.5 µm was employed. The forward segment of the collected force-displacement (*F-z*) curves was analysed using an open-source software (43). To convert *F-z* curves into force-indentation (*F-Z*) curves, the contact point was found using a goodness-of-fit algorithm (69). Subsequently, the Hertz model (**Equation 1**) was fitted up to an indentation of 10% of the probe radius (i.e., δ = 0.1*R*) to obtain *E*, assuming Poisson’s ratio (*v*) to be 0.5:

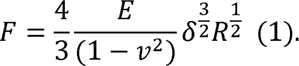

At least 3 hydrogels per condition were tested, with a minimum of 50 indentations per hydrogel. Each indentation was spaced at least twice the contact radius (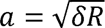) from the previous one to ensure testing a different point. All measurements were conducted at room temperature in Milli-Q water on thin hydrogel films immobilized on silanized coverslips.

### Determination of the stress relaxation

Stress relaxation measurements were conducted using a fiber-optic based nanoindentation device (Chiaro, Optics 11 Life) mounted on top of an inverted optical microscope (Axiovert 200M, Zeiss), adapting a previously described approach (18). The instrument operated in closed-loop mode (Indentation mode), maintaining a constant indentation depth, δ, over time to characterise the local stress relaxation response of the material. A cantilever with a *k* of 0.52 N/m and *R* of 27.5 µm was employed for all hydrogels. Hydrogels underwent indention at a strain rate of 30 µm/s until an indentation depth of 3 μm was reached (about 10% of the probe radius, *R*), maintained for 60 seconds. This corresponds to an applied strain (ε) of ∼7 % given (70):

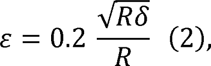

which fulfils the small strain approximation (70). Concurrently, the load signal was recorded. The acquired data were analysed using a previously described open source Jupyter notebook available on GitHub (18). At least two hydrogels per conditions were tested, with over 100 points per hydrogel, spaced at least twice the contact radius (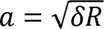) from the previous one to ensure different points were tested. All measurements were performed at room temperature in Milli-Q water on thin hydrogel films immobilized on silanized coverslips.

### Bulk rheological characterisation

Bulk rheology measurements were conducted using a Modular Compact Rheometer (MCR 302e, Anton Paar) equipped with a parallel plate geometry. A 15 mm diameter upper plate was employed for all tests. In brief, bulk gels were prepared by dispensing 400 µL of the pre-polymer solution on a PCTFE sheet (Agar Scientific), which was then covered with a hydrophobic glass coverslip (treated with RainX solution, RainX) of 18 mm in diameter. This process resulted in hydrogels of approximately 18mm in diameter with a thickness of ∼1 mm. The hydrogels were allowed to polymerize for 1 hour at room temperature. Subsequently, the coverslip-hydrogel-PCTFE sandwich was immersed in cold Milli-Q water for 1 hour, and hydrogels were gently detached. After washing the hydrogels three times in Milli-Q water, they were stored at 4°C overnight in Milli-Q water to facilitate swelling. Prior to measurements, hydrogels were punched to 15 mm to match the rheometer’s upper plate dimensions. The upper plate of the rheometer was brought into contact with the sample until a normal force of ≈0.1N was reached to ensure good contact without introducing compressional stiffening effects (71). Subsequently, an amplitude sweep between 0.01% and 1% strain (angular frequency of 10 rad/s) was performed to identify a suitable strain within the linear viscoelastic regime for subsequent frequency sweeps. A frequency sweep was then conducted at 1% strain between 0.0186 and 8.6100 Hz (0.1 to 54 rad/s). All measurements were performed at room temperature in Milli-Q water, and the hydration of the sampled was maintained using a solvent trap. Three independent samples per condition were tested.

### Cell culture

MCF-10A cells were culture in DMEM/F-12 (Thermo Fisher) supplemented with 5% Horse Serum (Thermo Fisher), 20 ng/mL epidermal growth factor (EGF) (Peprotech), 0.5 mg/mL Hydrocortisone (Merck), 100 ng/mL Cholera Toxin (Enzo Life Sciences), 10 µg/mL insulin from bovine pancreas (Merck) and 0.1% Penicillin/Streptomycin (Pen/Strep) (Merck). Cells were passaged every 3-4 days when reaching confluence and plated at a 1:4 dilution (∼2 million cells/T75 flask). For cell passaging, the culture media was aspirated, and cells were washed with 10 mL of 1x PBS. After aspirating the PBS, cells were incubated with 2mL of 0.25% trypsin (Merck) at 37 °C for 10 minutes. Trypsinisation was halted by adding 4 mL of media, and cells were spun at 1.3 rpm for 5 minutes in a 15 mL falcon tube to obtain a pellet. The old media was aspirated, and cells were resuspended in 1mL of fresh media before being plated. Cells were used up to a maximum of 15 passages.

### Cell plating on 2D substrates

Prior to protein functionalization and cell seeding, both PAAm hydrogels and glass coverslips (170 µm thick) underwent UV sterilization for 30 minutes. Cells were trypsinized following the cell culture protocol and then plated at 20,000 cells per gel (approximately 7,000 cells/mL or roughly 5,000 cells/cm^2^, considering a 22mm hydrogel surface). After plating, cells were allowed to adhere for 24 hours at 37°C with 5% CO_2_ before initiating any experiment.

### Cell migration experiments

MCF10-A cells were seeded following the procedure outlined above. To arrest cell division before to commencing time-lapse experiments, cell were treated with 5 µg/mL Mitomycin C (Merck) for 1 hour. Subsequently, Mitomycin C was removed through a single wash with fresh media, coverslips were transferred in an Attofluor^TM^ chamber (ThermoFisher), and fresh media was added before initiating imaging. Time-lapse experiments were conducted on either a Nikon C1 or a Nikon Ti2 A1R HD25 inverted microscope (Nikon, Japan) using either a 20x/0.45 or 20x/0.75 DIC objective, respectively. Images were taken at 3-minutes intervals over a 15-hour period. An incubation chamber was employed to maintain CO_2_ levels at 5% and temperature at 37°C during imaging. Multiple positions (N>5) were imaged in all cases using a motorized stage. Each condition underwent at least two independent experiments.

### Immunocytochemistry

Staining of samples on both hydrogels and glass coverslips was carried out following the same protocol. Samples were washed three times with PBS and fixed with 4 % Paraformaldehyde (PFA) in PBS for 12 minutes at room temperature either after cell migration experiments or at least after 24 hours of plating cells. Subsequently, samples were then washed three times with PBS, with the last wash lasting for 5 minutes, and permeabilised using 0.05% Triton X-100 in PBS for 10 minutes at room temperature. Following another round of three washes with PBS, samples were blocked with a solution of 5 v/v % Fetal Bovine Serum (FBS, Gibco) and 1 w/v % Bovine Serum Albumin (BSA, Merck) in PBS for 30 minutes at room temperature. After an additional three washes with PBS, a solution of 1% BSA in PBS containing the primary antibody, Alexa Fluor 488 Phalloidin 1:200 (Thermofisher) and DAPI 1:200 (Thermofisher) was added to the samples and incubated for 45 minutes at 37°C. We utilized either the monoclonal anti-vinculin antibody produced in mouse (Merck, Ref: V9131) at 1:200 or the YAP monoclonal antibody (M01 clone 2F12, Abnova, Ref: H00010413-M01) at 1:100 as primary antibodies. Subsequently, samples were washed three time with PBS, and incubated with goat anti-mouse Alexa Fluor 555 secondary antibody (Thermofisher) (dilution 1:200) or Cy3-conjugated Rabbit Anti Mouse (Jackson ImmunoResearch) (dilution 1:250) in a solution of 1% BSA in PBS for 45 minutes at 37°C. Following another round of three washes with PBS, samples were mounted using ProLong Diamond Antifade Mountant (Thermofisher) and stored at 4 °C until imaging.

### Image acquisition for fixed samples

Images of the actin cytoskeleton on 2D homogeneous substrates were captured using either a Nikon C1 inverted fluorescence microscope with a ×20/0.45NA Plan Fluor objective and ×60/1.4 Plan Apo oil immersion objective; or a Zeiss LSM 980 confocal microscope using a x20/0.8 Plan Apo or ×63/1.4 Plan Apo oil immersion objective. Vinculin images for homogeneous 2D substrates were acquired using a Nikon C1 inverted fluorescence microscope with a 60x/1.4 Plan Apo oil immersion objective. YAP images for homogeneous 2D or micropatterned 1D substrates were acquired using a Zeiss LSM 980 confocal microscope using a ×20/0.8 Plan Apo objective or ×63/1.4 Plan Apo oil immersion objective. For actin cytoskeleton images on 1D micropatterned substrates, a Nikon C1 or A1 Ti2 A1R HD25 inverted microscope (Nikon, Japan) with a ×60/1.4 Plan Apo objective or ×100/1.35 Plan Apo silicone immersion objective was used. Vinculin images on 1D micropatterned substrates were obtained using a Nikon A1 Ti2 A1R HD25 inverted microscope equipped with a 100x/1.35 Plan Apo silicone immersion objective.

### Actin flow experiments

MCF-10A cells were plated on PAAm hydrogels, following the previously described protocol for cell plating in 2D PAAm hydrogels. Prior to imaging, cells were incubated with SPY555-FastAct™ (Spirochrome) for 2 hours at 37°C following the manufacturer’s protocol (dilution 1:1000 in culture medium). The hydrogels were then transferred to an Attofluor^TM^ chamber (Thermofisher). Live actin imaging was performed on a Zeiss LSM 980 confocal microscope using a Plan Apo 40x/1.3 oil immersion objective. An incubation chamber was used to maintain CO_2_ levels at 5% and temperature at 37 °C throughout the imaging session. Images were acquired at a rate of one per second for a total duration of two minutes.

### Cell projected area and cell morphology

Cell projected area, circularity and aspect ratio were quantified in Fiji (72) by applying a Gaussian blur filter (sigma=2), followed by a default threshold to segment individual cells and quantify parameters of interest. The threshold was manually adjusted to capture the entire cell cytoskeleton.

### Focal adhesions analysis

Focal adhesions were analysed in Fiji (72) by following a previously published protocol (73). Briefly, images were cropped to include only one cell per image. Subsequently, background subtraction was performed using a sliding paraboloid and rolling ball radius of 50. Local contrast was then enhanced by applying CLAHE using block size=19, histogram bins=256, and maximum slope=6. The image underwent an exponential transformation to minimise background. Automatic adjustments were made to image brightness and contrast, followed by the application of a Log3D filter with sigmax=sigmay=3 applied. The LUT of the image was inverted, and automatic threshold was applied to convert the image to a binary. A watershed algorithm was then used to eliminate incorrectly clustered adhesions. Finally, the “Analyse Particles” command was executed (minimum 20 pixels to maximum infinity) to quantify focal adhesions area and count per cell. Alternatively, for 1D micropatterned hydrogels, images were inspected to count the number of cells having front/rear vinculin patches.

### Nuc/Cyto YAP ratio

Nuclear (Nuc) to Cytoplasmic (Cyto) ratio was quantified using Fiji (72). Initially, nuclear area (*A*_nuc_) was calculated by applying a gaussian blur (sigma=2) and segmenting nuclei (DAPI channel) via the default threshold. Subsequently, cell area (*A*_cell_) was calculated by applying a gaussian blur (sigma=2) and segmenting the actin cytoskeleton (Phalloidin channel) via the Default threshold. Nuclear and cellular outlines were saved in the Fiji ROI manager and utilized to quantify integrated density of the cell (*YAP*_cell_) and the nucleus (*YAP*_nuc_) in the YAP channel. Therefore, Nuc/Cyto YAP ratio was calculated as follows:

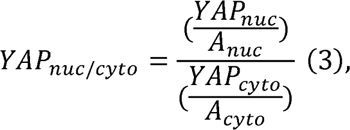

Where

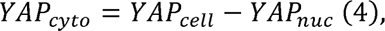

And

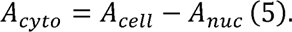

### Cell tracking and migration analysis

DIC time-lapses were imported in Fiji (72), converted to 8-bit format, and analysed using CellTracker (74). In brief, manual tracking was performed by clicking on the cell’s centroid every two frames, and linear interpolation was applied to compute their position over time. Only cells that migrated a distance greater than three cell bodies were considered migratory and therefore tracked. Cell trajectories were aligned to a common origin and are shown for a tracking time of 5 hours. Cell speed was computed as the mean instantaneous speed considering the entire tracking period. For a tracking time of 5 hours, MSD, directionality ratio, and autocorrelation were computed using a previously published protocol (61). Average MSD curves were fitted according to:

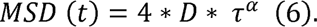

In Equation (6), α represents the anomalous diffusion exponent describing the type of diffusion of the system (α = 1 for a diffusive system, α > 1 for a super diffusive system and α < 1 for a sub diffusive system), τ is the lag time and *D* is the diffusion coefficient (19, 75).

### Retrograde actin Flow

To determine the retrograde actin flow at the cell leading edge (lamellipodium), time-lapses were initially converted to 8-bit format, underwent background subtraction, contrast enhanced, and were subjected to a Gaussian blur filter (sigma=1.5). Subsequently, kymographs were generated using the Multi Kymograph plugin in Fiji with a line width of 1. Actin retrograde flow speed was then computed using the bounding rectangle parameters as follows:

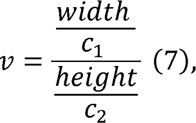

Where *width* is the width of the bounding rectangle in pixels, *c_1_* is the spatial conversion factor (px/μm), *height* is the height of the bounding rectangle in pixels, and *c_2_* is the temporal conversion factor (px/s). This calculation yields the actin retrograde flow speed (*v*) in units of µm/s, which is subsequently converted to nm/s.

### Statistical analysis

All statistical analyses were performed in Prism v10 (GraphPad). Two-tailed unpaired student t tests were employed when comparing two conditions. For comparisons involving more than two conditions with variations in two variables, two-way ANOVAS with Bonferroni’s or Tukey’s multiple comparisons tests were performed. For comparisons involving more than two conditions with variations in three variables, three-way ANOVAS were performed with Tukey’s multiple comparisons test. Specific tests conducted for each analysis are detailed in the respective figure captions. Comparisons were considered significant when the p value was at least smaller than 0.05.

## Authors contribution

S.G., G.C., M.S.-S. conceived the project. S.G. and M.S.-S. and M.V. supervised the project. G.C. performed all experiments and analysed all data. M.A.G.O. performed nanoindentation experiments together with G.C. G.C. wrote the first original draft of the article, which was reviewed and edited by S.G. The article was read and corrected by all authors, who contributed to the interpretation of results. Funding was acquired by M.V., M.S.-S. and S.G.

## Conflict of interest

The authors declare no conflicts of interest.

## Acknowledgements

G.C. acknowledges all members of the CeMi (University of Glasgow) and Mechanobiology and Biomaterials group (University of Mons) for helpful advice and discussion. G.C. acknowledges financial support from the UKRI for PhD funding, the Research Institute for Biosciences (UMONS) for the invitation grant and EMBO for the scientific exchange grant # 10122 that made this collaborative work possible. S.G. acknowledges funding from FEDER Prostem Research Project no. 1510614 (Wallonia DG06), the F.R.S.-FNRS Epiforce Project no. T.0092.21, the F.R.S.-FNRS CellSqueezer Project no. J.0061.23, the F.R.S.-FNRS Optopattern Project no. U.NO26.22 and the Interreg MAT(T)ISSE project, which is financially supported by Interreg France-Wallonie-Vlaanderen (Fonds Européen de Développement Régional, FEDER-ERDF), Programme Wallon d’Investissement Région Wallone pour les instruments d’imagerie (INSTIMAG UMONS #1910169). M.S-S. is grateful for financial support from the European Research Council AdG (Devise, 101054728). IBEC is member of CERCA Programme / Generalitat de Catalunya. M.C. acknowledges funding from MRC (MR/S005412/1) and Royal Society (RGS/R1/231400).

## Data Availability

Data will be shared from the corresponding author upon reasonable request.

## Supporting information

**Supplementary Table S1.**
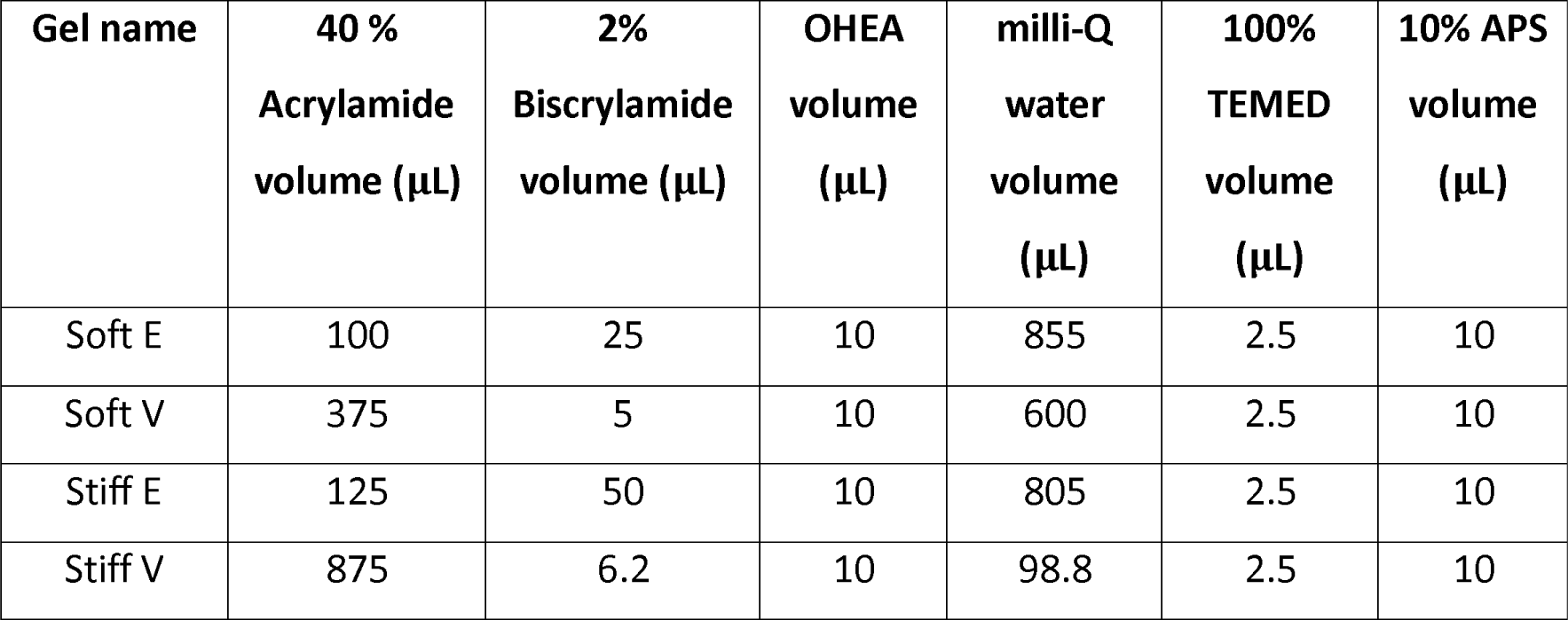
Hydrogels’ formulations used in this study. E (elastic), V (viscoelastic). OHEA = oxidised N-hydroxyethyl acrylamide, TEMED = tetramethylethylenediamine, APS = ammonium persulfate.

**Supplementary Figure 1–.**
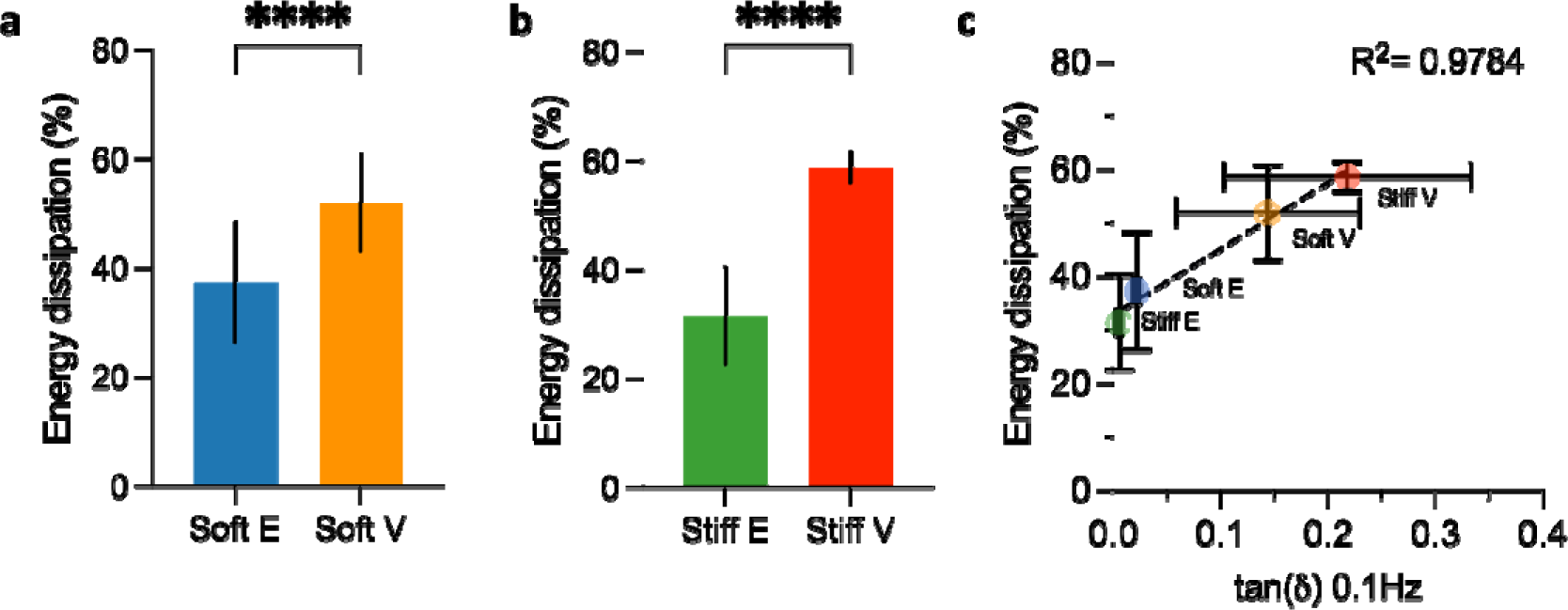
Viscoelastic properties of soft and stiff polyacrylamide hydrogels. **(a)** Average energy dissipation over ∼60 s for soft elastic (Soft E) and viscoelastic (Soft V) hydrogels. Data is shown as mean ± standard deviation (n = 131 for Soft E, n = 151 for soft V over at least two independent samples). **(b)** Average energy dissipation over ∼60 s for stiff elastic (Stiff E) and viscoelastic (Stiff V) hydrogels. Data is shown as mean ± standard deviation (n = 142 for stiff E, n = 121 for stiff V over at least two independent samples). **(c)** Average energy dissipation over ∼60 plotted against the tan(δ) at 0.1 Hz. Data is shown as mean ± standard deviation, where the number of points for the relaxation half time is the same as in a-b, and the number of independent samples for the tan(δ) is 3 (R^2^=0.9784).

**Supplementary Figure 2–.**
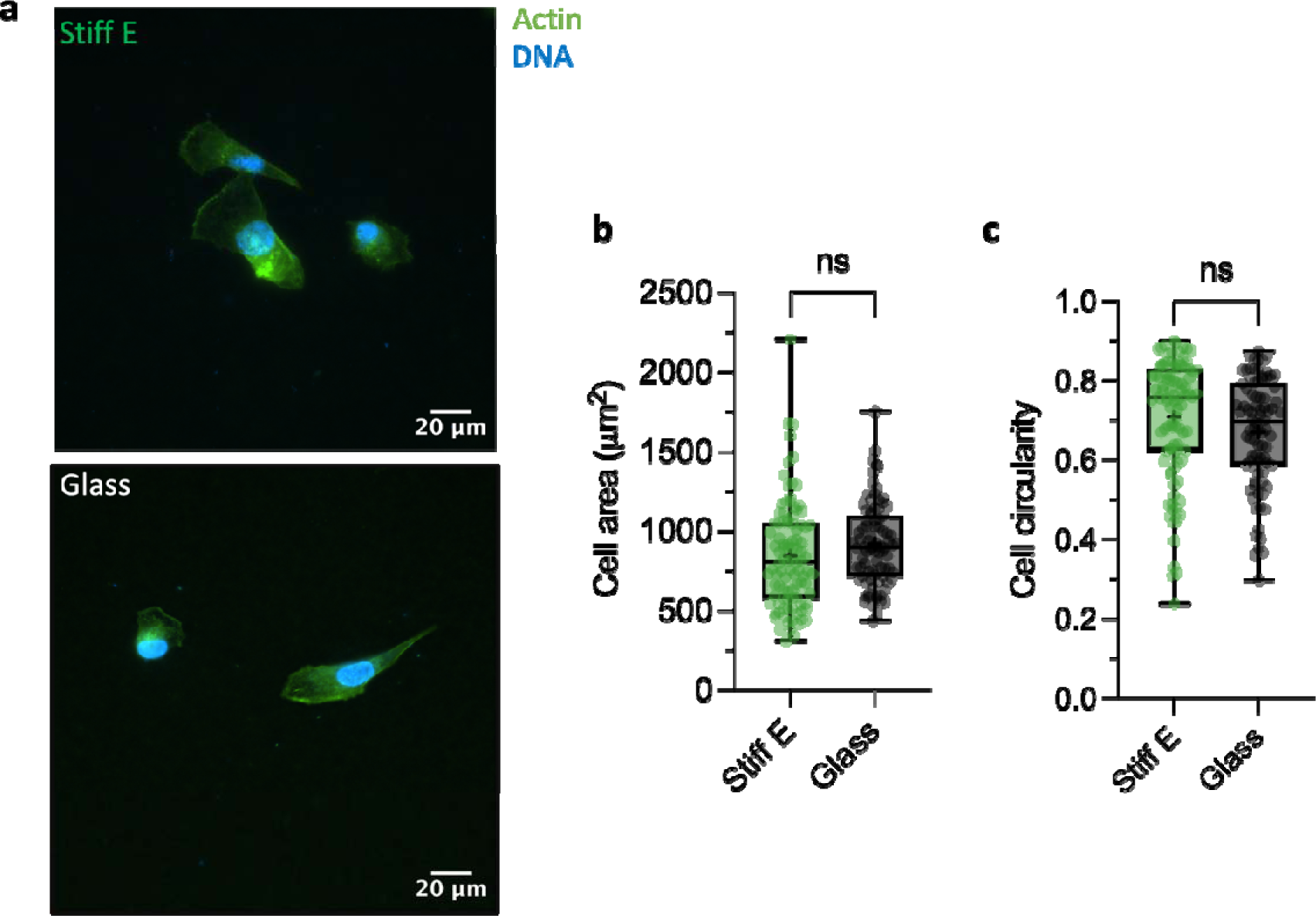
Cell spreading area and circularity are the same on stiff elastic (Stiff E) hydrogels and fibronectin coated glass coverslips (Glass). **(a)** Representative Actin/DNA images of MCF-10A cells cultured on stiff elastic hydrogels (Stiff E) and fibronectin-coated glass coverslips. **(b)** Quantification of MCF-10A cell spreading area on stiff elastic (Stiff E) hydrogels (n = 100 cells) and fibronectin-coated glass coverslips (Glass) (n = 86 cells) from at least two independent experiments. ns p = 0.0985, two-tailed unpaired t-test. **(c)** Quantification of MCF-10A cell circularity on stiff elastic (Stiff E) hydrogels (n = 100 cells) and fibronectin-coated glass coverslips (Glass) (n = 86 cells) from at least two independent experiments. ns p = 0.0810, two-tailed unpaired t-test.

**Supplementary Figure 3–.**
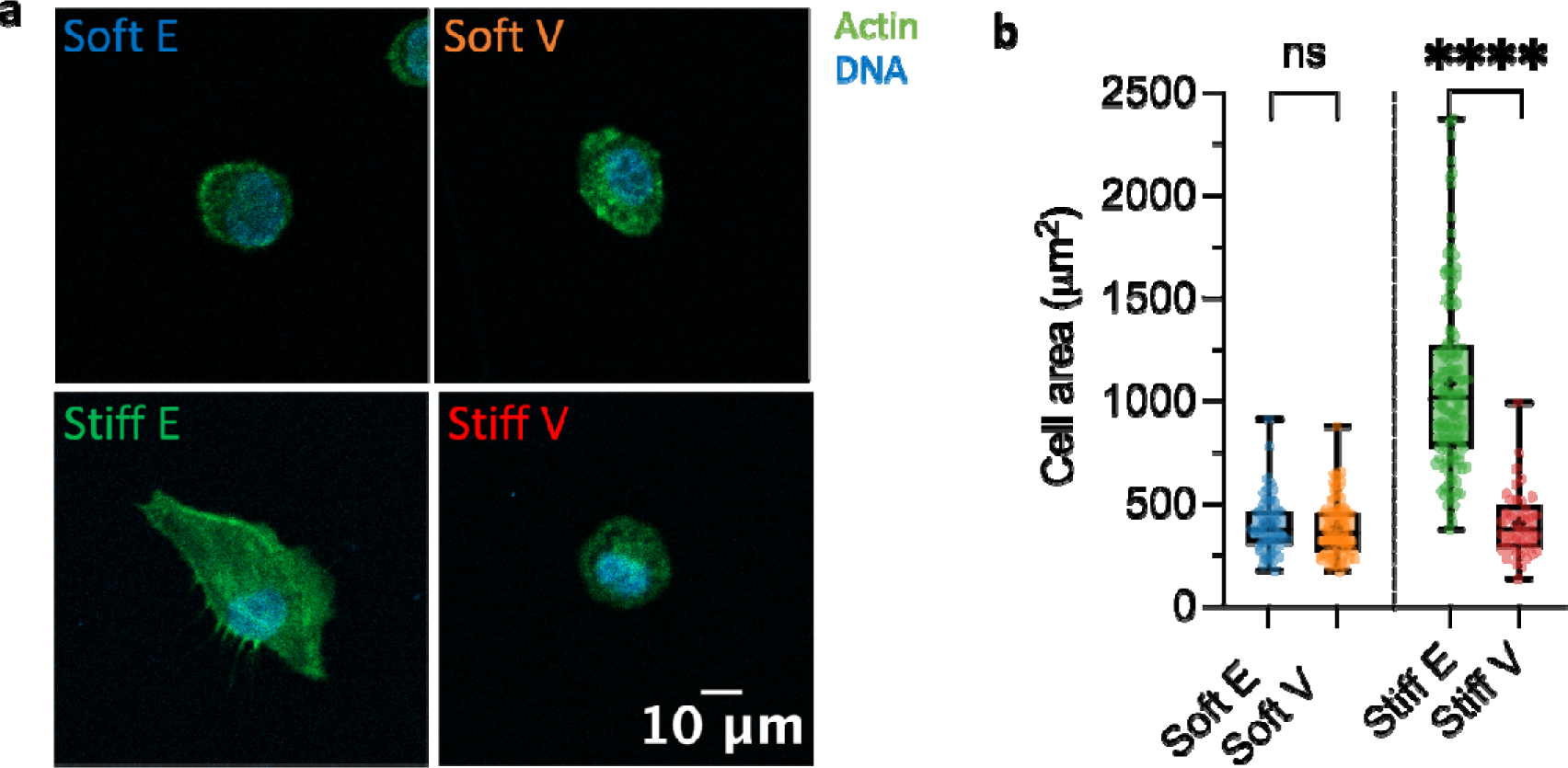
Cell spreading on elastic and viscoelastic polyacrylamide hydrogels coated with low fibronectin concentration. (a) Representative Actin/DNA images of MCF-10A cells cultured on viscoelastic polyacrylamide hydrogels coated with low fibronectin concentration. (b) Quantification of the cell spreading area of MCF-10A cells cultured on viscoelastic polyacrylamide hydrogels coated with low fibronectin concentration (n= 61 cells for Soft E, n = 84 cells for Soft V, n = 138 cells for Stiff E, n = 41 cells for Stiff V from two independent experiments). ns p >0.9999, ****p<0.0001, two-way ANOVA with Bonferroni’s multiple comparisons test.

**Supplementary Figure 4–.**
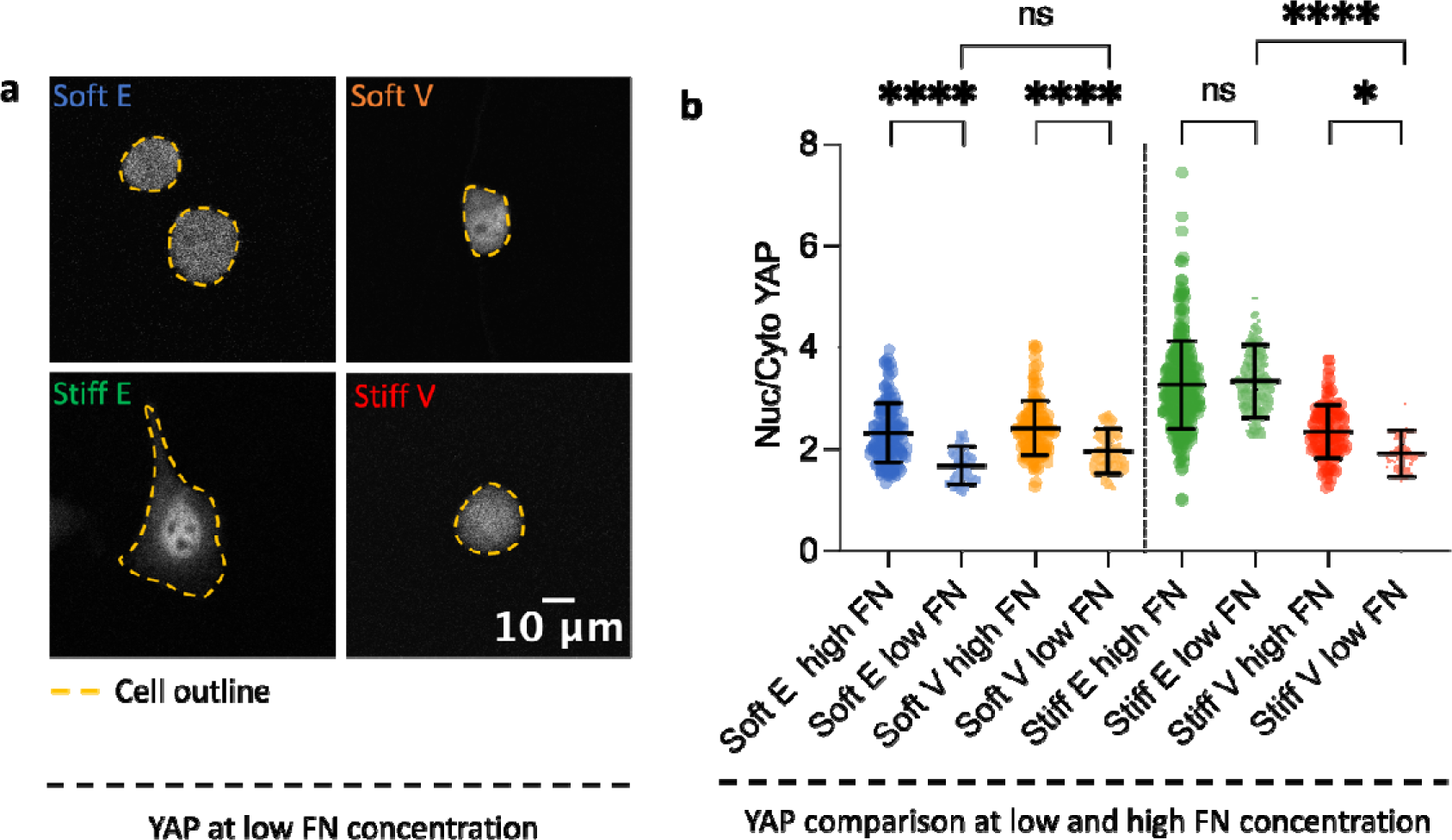
YAP nuclear translocation on elastic and viscoelastic polyacrylamide hydrogels coated with low FN concentration. (a) Representative fluorescent images of MCF-10A cells cultured on viscoelastic polyacrylamide hydrogels coated with low FN concentration and stained for YAP. The cellular outline is depicted by a dashed yellow line. (b) Quantification of the Nuclear to Cytoplasmic (Nuc/Cyto) YAP ratio of MCF-10A cells cultured on viscoelastic polyacrylamide hydrogels coated with high and low FN concentration. Data for high FN concentration is presented in Fig. 2. Data for low FN concentration is as follows: n= 61 cells for Soft E, n = 84 cells for Soft V, n = 138 cells for Stiff E, and n = 41 cells for Stiff V from two independent experiments. Data is shown as mean ± SD. ns p > 0.05, * p < 0.0107, **** p< 0.0001, three-way ANOVA with Tukey’s multiple comparisons test.

**Supplementary Figure 5–.**
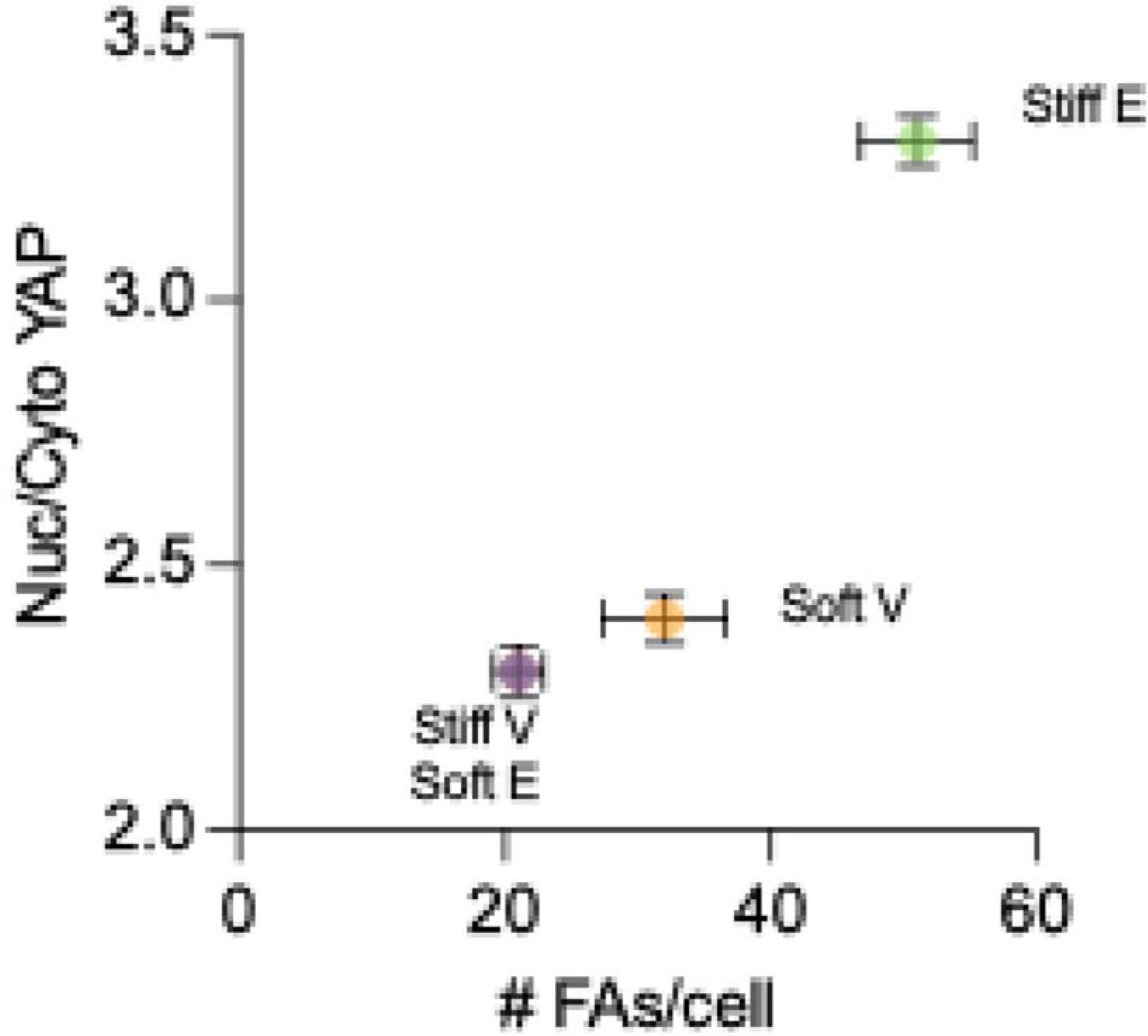
Evolution of the Nuclear to Cytoplasmic YAP ratio (Nuc/Cyto) YAP as a function of the number of focal adhesions per cell (# FAs/cell) on elastic and viscoelastic polyacrylamide hydrogels. Data is shown as mean ± SEM. For YAP, sample size is as follows: soft E: n=157 cells, Soft V: n=128 cells, Stiff E: n=320 cells, and Stiff V: n=113 cells, from at least two independent experiments. For FAs, sample size is as follows: soft E: n = 33 cells, soft V: n=41 cells, Stiff E: n=58 cells, and stiff V: n=27 cells, from at least two independent experiments.

**Supplementary Figure 6–.**
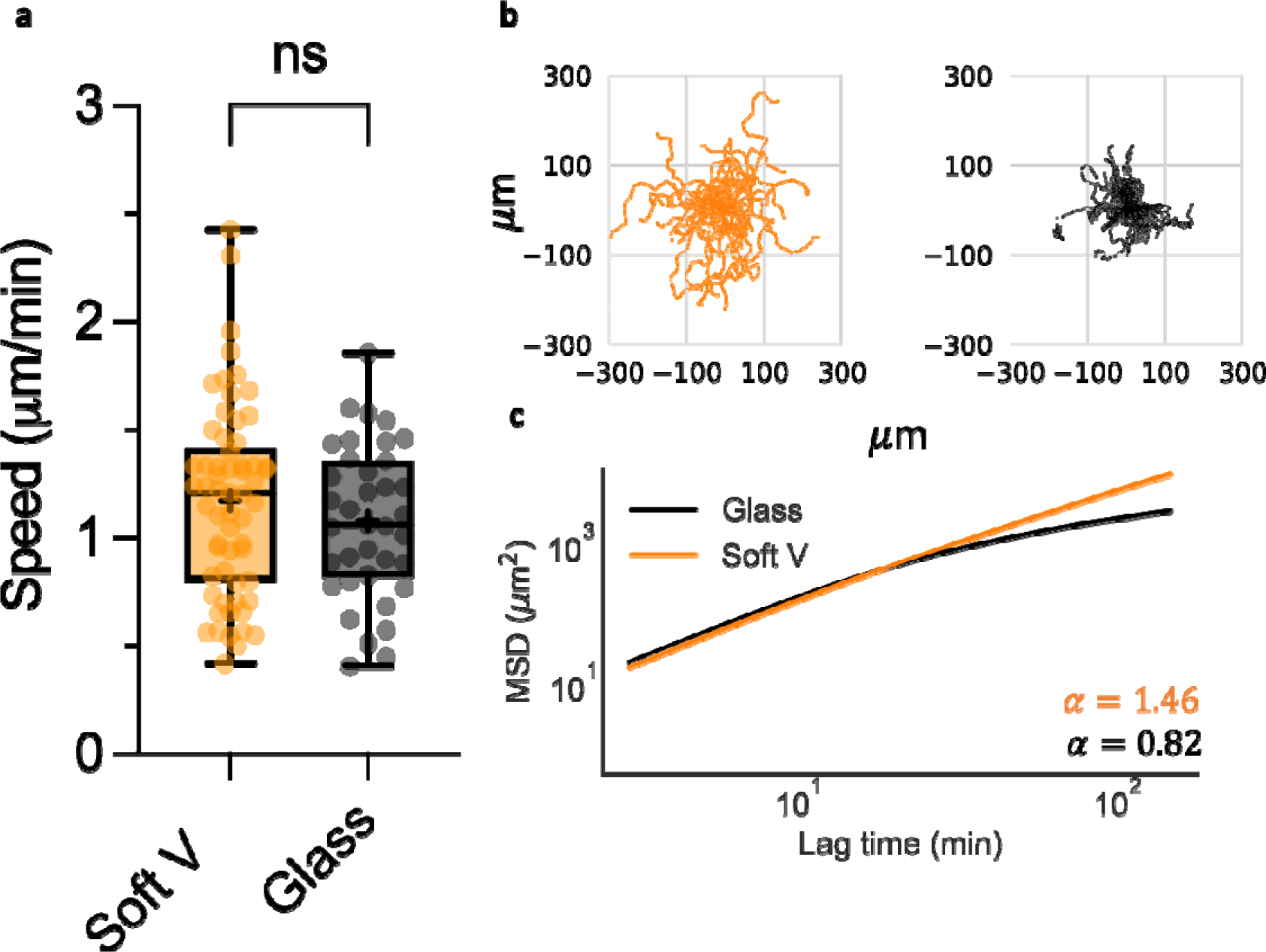
MCF-10A cells exhibit comparable migration speed but higher directional persistence on soft viscoelastic (Soft V) matrices in contrast to fibronectin-coated glass coverslips (Glass). **(a)** Quantification of MCF-10A cell migration speed (n= 61 cells for Soft V, n = 37 cells for Glass) from at least two independent experiments. ns p = 0.2842, two-tailed unpaired t-test. **(b)** Representative trajectories of MCF-10A cells migration on soft V and glass (n=43 cells for Soft V, n= 33 cells for Glass) from at least two independent experiments. **(c)** Average mean square displacement (MSD) vs lag-time for MCF-10A cells on Soft V and Glass from at least two independent experiments. The diffusion exponent, α, is shown in the graph. Data is shown as mean ± SEM (n=43 cells for Soft V, n= 33 cells for Glass).

**Supplementary Figure 7–.**
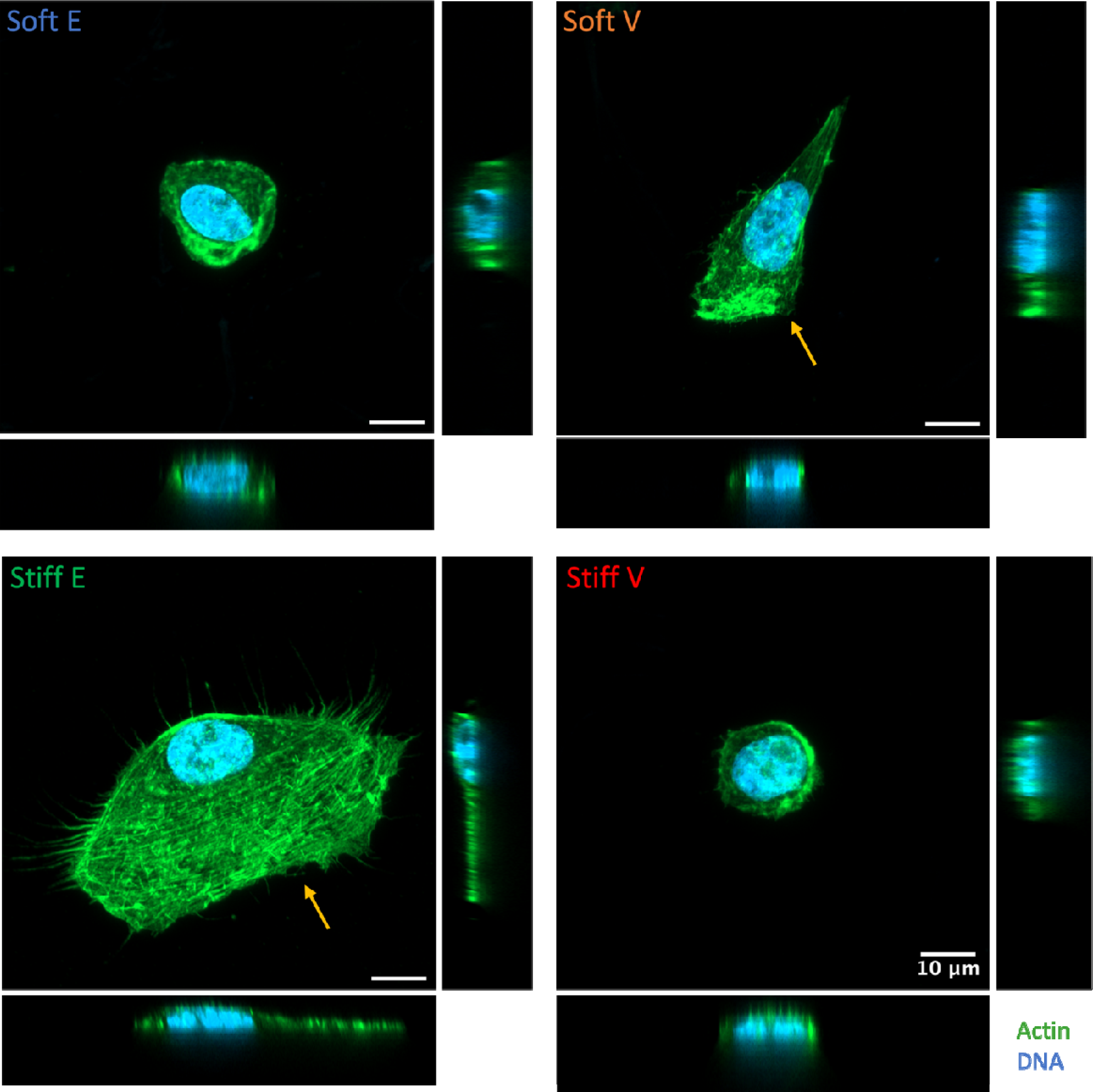
Lamellipodium formation on elastic and viscoelastic polyacrylamide hydrogels. Representative maximum intensity projections of MCF-10A cells cultured on elastic and viscoelastic polyacrylamide hydrogels. Corresponding (*xz*) projections (horizontal) and (*yz*) projections (vertical) are shown alongside each image. Lamellipodium is indicated by a yellow arrow. Images for soft E and stiff V are the same as in Fig. 1a. All scale bars are 10 µm.

**Supplementary Figure 8–.**
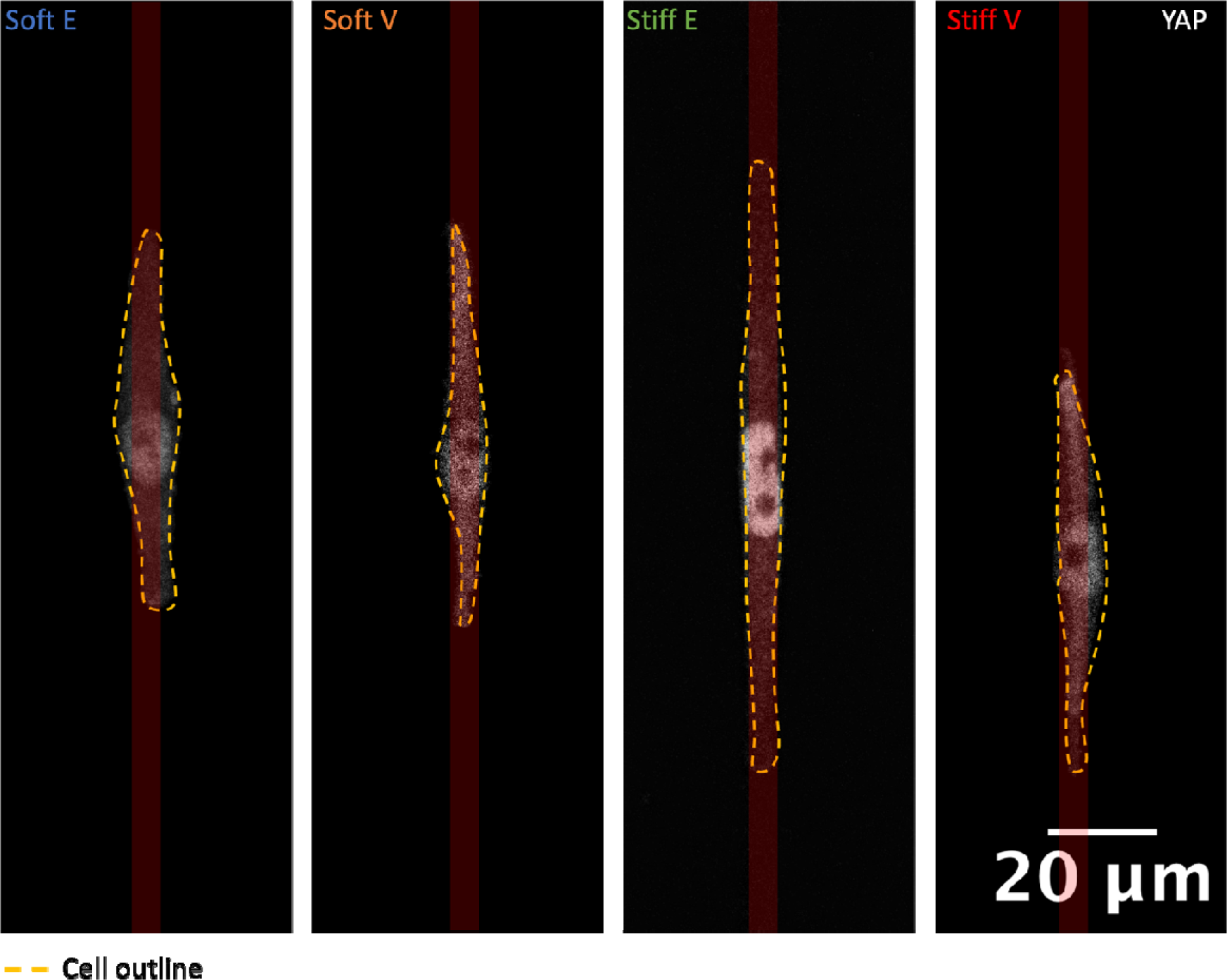
Representative images of YAP nuclear translocation of MCF-10A cells on 5 μm micropatterned fibronectin lines on elastic and viscoelastic polyacrylamide hydrogels. Fibronectin lines are schematically represented in red for clarity. The cellular outline is depicted by a dashed yellow line.

**Supporting Movie S1 –** Time lapse movie in DIC mode of MCF-10A cells migrating for 15h on a stiff elastic (Stiff E) hydrogel substrate.

**Supporting Movie S2 –** Time lapse movie in DIC mode of MCF-10A cells migrating for 15h on a stiff viscoelastic (Stiff V) hydrogel substrate.

**Supporting Movie S3 –** Time lapse movie in DIC mode of MCF-10A cells migrating for 15h on a soft elastic (Soft E) hydrogel substrate.

**Supporting Movie S4 –** Time lapse movie in DIC mode of MCF-10A cells migrating for 15h on a soft viscoelastic (Soft V) hydrogel substrate.

**Supporting Movie S5 –** Confocal volume rendering of a single MCF-10A cell on a stiff elastic (Stiff E) hydrogel substrate stained for actin (green) and DNA (blue).

**Supporting Movie S6 –** Confocal volume rendering of a single MCF-10A cell on a stiff viscoelastic (Stiff V) hydrogel substrate stained for actin (green) and DNA (blue).

**Supporting Movie S7 –** Confocal volume rendering of a single MCF-10A cell on a soft elastic (Soft E) hydrogel substrate stained for actin (green) and DNA (blue).

**Supporting Movie S8 –** Confocal volume rendering of a single MCF-10A cell on a soft viscoelastic (Soft V) hydrogel substrate stained for actin (green) and DNA (blue).

**Supporting Movie S9 –** Time lapse movie in confocal fluorescence mode of a single MCF-10A tagged with Spy-555-FastAct on a stiff elastic (Stiff E) hydrogel substrate.

**Supporting Movie S10 –** Time lapse movie in confocal fluorescence mode of a single MCF-10A tagged with Spy-555-FastAct on a stiff viscoelastic (Stiff V) hydrogel substrate.

**Supporting Movie S11 –** Time lapse movie in confocal fluorescence mode of a single MCF-10A cell tagged with Spy-555-FastAct on a soft elastic (Soft E) hydrogel substrate.

**Supporting Movie S12 –** Time lapse movie in confocal fluorescence mode of a single MCF-10A cell tagged with Spy-555-FastAct on a soft elastic (Soft E) hydrogel substrate.

**Supporting Movie S13 –** Time lapse movie of MCF-10A cells migrating for 15h on a soft elastic (Soft E) micropatterned hydrogel substrate with 5 μm fibronectin lines (DIC mode in grey and DNA in blue).

**Supporting Movie S14 –** Time lapse movie of MCF-10A cells migrating for 15h on a soft viscoelastic (Soft V) micropatterned hydrogel substrate with 5 μm fibronectin lines (DIC mode in grey and DNA in blue).

**Supporting Movie S15 –** Time lapse movie of MCF-10A cells migrating for 15h on a stiff elastic (Stiff E) micropatterned hydrogel substrate with 5 μm fibronectin lines (DIC mode in grey and DNA in blue).

**Supporting Movie S16 –** Time lapse movie of MCF-10A cells migrating for 15h on a stiff viscoelastic (Stiff V) micropatterned hydrogel substrate with 5 μm fibronectin lines (DIC mode in grey and DNA in blue).

## Notes

### Competing Interest Statement

The authors have declared no competing interest.

